# Metagenomics of the MAST-3 stramenopile, *Incisomonas,* and its associated microbiome reveals unexpected metabolic attributes and extensive nutrient dependencies

**DOI:** 10.1101/2025.02.28.640650

**Authors:** Dominic E Absolon, Victoria LN Jackson, Adam Monier, Alison G Smith, Katherine E Helliwell

**Affiliations:** Department of Plant Sciences, University of Cambridge, Cambridge, UK; Biosciences, Faculty of Health and Life Sciences, University of Exeter, Exeter, EX4 4QD, UK; Living Systems Institute, University of Exeter, Exeter, EX4 4QD, UK; Marine Biological Association, Citadel Hill, Plymouth, PL1 2PB, UK

## Abstract

Protists are polyphyletic singled-celled eukaryotes that underpin global ecosystem functioning, particularly in the oceans. Most remain uncultured, limiting investigation of their physiology and cell biology. <u>MA</u>rine <u>ST</u>ramenopiles (MASTs) are heterotrophic protists that, although related to well-characterised photosynthetic diatoms and parasitic oomycetes, are poorly studied. The Nanomonadea (MAST-3) species *Incisomonas marina* has been maintained in co-culture with a bacterial consortium, offering opportunities to investigate the metabolic attributes and nutritional dependencies of the community. Employing a metagenomics approach, the 68 Mbp haploid genome of *I. marina* was retrieved to an estimated completeness of 93%, representing the most complete MAST genome so far. We also characterised the diversity of, and assembled genomes for, 23 co-cultured bacteria. Auxotrophy of *I. marina* for B vitamins (B_1_, B_2_, B_6_, B_7_ and B_12_), but not vitamins C, B_3_, B_5_ and B_9_ was predicted. Several bacteria also lacked complete B-vitamin biosynthesis pathways, suggesting that vitamins and/or their precursors are exchanged in the consortium. Moreover, *I. marina* lacked the ability to synthesise half the protein amino acids, although genes encoding the complete urea cycle were identified, like diatoms; this may play a role in recycling organic nitrogen compounds. Unexpectedly, we also identified the gene *DSYB* for dimethylsulphoniopropionate (DMSP) biosynthesis. Biosynthesis of this important stress-protectant and bacterial chemoattractant is typically found in photosynthetic eukaryotes and has not before been identified in heterotrophic stramenopiles. Together, our study reveals the metabolic attributes of a hitherto understudied organism, advancing knowledge of the evolution and adaptations of the stramenopiles and informing future culturing efforts.

## Introduction

Marine protists play crucial roles in nutrient and carbon cycling in the oceans (Worden et al. 2015). However, the majority of heterotrophic eukaryotic microorganisms remain uncharacterised, their existence evident only through molecular surveys using DNA metabarcoding approaches (del Campo et al. 2014). A major polyphyletic assemblage described within this unexplored protistan diversity is the <u>MA</u>rine <u>ST</u>ramenopiles, so-called ‘MASTs’ (Massana et al. 2004). An emerging picture from environmental surveys (e.g. *Tara* Oceans (Seeleuthner et al. 2018)) is that MASTs are abundant and globally distributed across marine habitats (Orsi et al. 2011; Logares et al. 2012; Logares et al. 2014; Massana et al. 2014). MASTs occupy several independent clades at the base of the stramenopile lineage, which encompasses a diverse range of eukaryotes. These include well-studied groups such as the heterotrophic oomycetes, a clade containing many parasitic species (Massana et al. 2006) and thraustochytrids, of biotechnological importance as a source of omega-3 fatty acids (Lyu et al. 2021). Moreover, there are many globally significant photosynthetic members, including diatoms and brown macroalgae; the diatoms alone are estimated to be responsible for ∼20% of planetary carbon fixation (Armbrust 2009).

In contrast, the different lifestyles and evolutionary origin of MAST species are poorly understood. Twenty-one distinct MAST clades have been described based on 18S rDNA analysis (Massana et al. 2014; Obiol et al. 2024) and they reportedly exhibit a range of different ecological strategies, including symbioses with diatoms and bacterial grazing (bacteriovory) (Massana et al. 2006; Gomez et al. 2016). Knowledge of the fundamental biology of MASTs has been hindered considerably by cultivation difficulties, likely due to their nutritional requirements, including reliance on other microbes in their environment (del Campo et al. 2014). Currently, just two MAST species have been cultured: *Incisomonas marina* (from the MAST-3 clade) (Cavalier-Smith and Scoble 2013) and *Pseudophyllomitus vesiculosus* (MAST-6) (Shiratori et al. 2017), both with associated microbiota. Currently, *I. marina* is the only publicly available strain. Moreover, genome sequencing efforts have generally relied upon single-cell genomics (SAGs) and metagenome assembled genomes (MAGs), recovered from environmental sampling approaches (Seeleuthner et al. 2018; Labarre et al. 2021). A major challenge of such techniques is poor genome recovery, with genome completeness ranging from 83% to just 7% (Labarre et al. 2021; Latorre et al. 2021). These studies have yielded details of some of the functional repertoire of MASTs, including aspects relating to motility, the phagocytosis apparatus, carbohydrate-active (CaZY) enzymes and rhodopsin encoding genes (Seeleuthner et al. 2018; Labarre et al. 2021; Latorre et al. 2021), and phylogenomic analysis of *I. marina* has been important in resolving aspects of stramenopile phylogeny (Derelle et al. 2016). However, while these studies have examined the presence of particular gene families, assessing key metabolic capabilities, such as energy utilisation and storage pathways, or biosynthesis of amino acids and cofactors that would indicate their nutritional needs and ecological lifestyle, is more challenging.

The diversity and wide array of ecological lifestyles represented by the stramenopiles makes this group ideal to study evolutionary transitions, including from heterotrophy and/or mixotrophy to autotrophy (Jirsová and Wideman 2024), and indeed back again (Kamikawa et al. 2022). Photosynthesis arose within the stramenopiles through endosymbiotic acquisition of a red algal derived plastid, driving genetic mixing between the protistan host and algal endosymbiont (Armbrust et al. 2004; Dorrell and Smith 2011). The combination of metabolic attributes likely contributed to the ecological success of the diatoms, as in other major marine algal taxa such as dinoflagellates and haptophytes (Sibbald and Archibald 2020). We now know that despite being photoautotrophs, diatoms take up significant amounts of organic substrates (Meyer et al. 2022), although whether this capacity has been shaped by their heterotrophic ancestry requires further exploration. Just as organic substrate usage may not be accurately predicted by inferred nutritional mode, the capacity for biosynthesis of specific metabolites need not be either. Many obligate heterotrophs (e.g. fungi and bacteria) often synthesise vitamins (Perli et al. 2020; Sultana et al. 2023), yet the majority algae are known to be auxotrophic for at least one B vitamin despite being photoautotrophic (Croft et al. 2006). These compounds have emerged as important currencies mediating microbial eukaryote-bacteria interactions (Croft et al. 2005; Durham et al. 2015; Cooper et al. 2019). B vitamins have even been predicted to flow from a bacterivorous protist (choanoflagellate) to bacterial associates, challenging typical models of trophic interactions in marine food webs (Needham et al. 2022). Similarly, amino acids and signalling molecules like dimethylsulfoniopropionate (DMSP) (Seymour et al. 2010; Kuhlisch et al. 2023), made by many photosynthetic stramenopiles (Curson et al. 2017; Wang et al. 2024), are widely exchanged. However, the biosynthetic capabilities of most MASTs are unknown, limiting understanding of their role in the broader marine ecosystems. Establishing the nutritional needs of MASTs is also crucial to facilitate culturing efforts.

Here, we took advantage of the publicly available *I. marina* co-culture, and characterised it at the physiological, morphological and genome sequence level, allowing detailed examination of the specific nutritional dependencies of *I. marina* and the co-cultured bacterial consortium, together with the identification of novel gene functions. This work helps establish *I. marina* as a model heterotrophic stramenopile with robust sequencing resources.

## Materials and Methods

### Culturing of *I. marina*

*I. marina* was acquired from the Culture Collection of Algae and Protozoa (culture number CCAP 997/1). The culture was maintained in artificial seawater for protozoa (ASWP; https://www.ccap.ac.uk/index.php/media-recipes/), together with a single barley grain as a carbon source, which was sterilised by boiling (10 minutes) before addition. Cultures were routinely maintained at 15 °C in a 12:12 light:dark cycle, sub-culturing every 16 weeks (as recommended by CCAP). For experimental work, cultures were transferred more frequently (every 2 weeks) to ensure cells were actively growing.

### Bacterial isolation and colony PCR

For the initial analysis of bacteria present in *I. marina* cultures, samples were spread on Marine broth (Difco) agar plates and incubated in the conditions described above. Colony PCR was performed on colonies with distinct morphological appearances to amplify 16S rRNA (V6-V8) sequences, using: ACGCGHNRAACCTTACC (forward) and ACGGGCRGTGWGTRCAA (reverse) primers. PCR was carried out using Red Taq kit (Sigma-Aldrich) and the cycling conditions were 95°C for 1 min followed by 30 cycles of 95°C for 0.5 min, 50 °C for 0.5 min and 72°C for 1.5 minutes, followed by a final extension at 70°C for 3 mins. PCR products were analysed using a 2% agarose gel. Illustra GFX Gel Band Purification kit was used to purify PCR products and subsequently send for Sanger sequencing. Resulting sequences were analysed in Geneious Prime and SINA taxonomy identification tool (Pruesse et al. 2012).

### Scanning Electron Microscopy (SEM)

Cells were grown on Melinex plastic coverslips (Agar Scientific) for 14 days, then briefly dipped twice in cold, de-ionised water to remove any buffer salts and quickly plunge-frozen in liquid nitrogen-cooled ethane. Samples were transferred to liquid nitrogen-cooled brass inserts and freeze-dried overnight in a liquid nitrogen-cooled turbo freeze-drier (Quorum K775X). Samples were mounted on aluminium SEM stubs using conductive silver paint (Agar Scientific) and coated with 15 nm iridium using a Quorum K575X sputter coater. Samples were viewed using a FEI Verios 460 scanning electron microscope run at 2.00 keV and 50 pA probe current. Secondary electron images were acquired using either an Everhard-Thornley detector in field-free mode (low resolution) or a Through-Lens detector in full immersion mode (high resolution).

### Metagenomics of *I. marina* culture

#### DNA extraction

*I. marina* cells (grown in six 200 ml cultures) were concentrated via centrifugation, resulting in a pellet consisting of a brown top layer and a lower white layer. Microscopy confirmed *I. marina* cells to be in the brown fraction. To reduce the bacterial load, which might interfere with sequence acquisition, these two fractions were manually separated, and DNA extracted from the brown fraction using a phenol-chloroform extraction. Biomass was resuspended in 400 µL of nuclease free water, and 400 µL of SDS-elution buffer (2% SDS, 100 mM Tris-HCl pH 8.0, 400 mM NaCl, and 40 mM EDTA, pH 8) added. The resulting sample was vortexed, before 800 µl of phenol:chloroform with iso-amyl alcohol (25:24:1 v/v) was added. The liquid suspension was vortexed and then centrifuged for 5 mins at 14,000 g, and the top phase transferred to a 2 mL Eppendorf tube. This was repeated twice before addition of 800 µL of chloroform:iso-amyl (24:1 v/v). The sample was then vortexed for 2 mins, before being centrifuged for 5 mins at 14,000 g and the top phase transferred to a new 2 ml Eppendorf tube. To this 2.5 × volume (∼1.4 ml) of (−20°C) ethanol 100% was added, and the sample incubated overnight at −20°C. The sample was then centrifuged for 30 mins at 16,000 g at 4°C, and the supernatant discarded. The pellet was resuspended in 400 µl of TE buffer, 1 µL of RNAse A added, and incubated for 1 h at 37 °C before one further phenol-chloroform extraction and ethanol precipitation. DNA was quality checked using a Nanodrop (ThermoFisher) and Qubit dsDNA HS assay kit (ThermoFisher).

### Long-read sequencing, quality control and genome assembly

The sequencing library was prepared using the Oxford Nanopore Technologies (ONT) ligation sequencing kit (SQK-LSK109) and the NEBNext FFPE DNA repair mix (M6630; New England Biolabs), NEBNext Ultra II end repair/dA-tailing module (E7546; NEB) and NEBNext quick ligation module (E6056; NEB), following the ONT protocol (version GDE_9063_V109_REVV_14AUG2019) with a modified incubation time for the repair and end-prep reaction. Briefly, DNA repair and end-prep reagents were added to 1 µg DNA and incubated for 20 min at 20 °C, followed by a final incubation step of 5 mins at 65 °C. Adapter ligation and clean-up were carried out according to the ONT protocol using long fragment buffer (LFB), and the eluted library was incubated at 37 °C for 10 min to improve the recovery of long fragments. Approximately 800 ng of DNA library was loaded onto a MinION SpotON flow cell (R9.4.1) and sequencing was carried out over 72 h. Base calling was performed with Guppy (ONT) in high-accuracy mode. The quality of the long-read data was assessed with LongQC (1.2.0) (Fukasawa et al. 2020) and nanoplot (1.34.0) (De Coster et al. 2018), both run with default parameters. Trimmomatic (0.39) (Bolger et al. 2014) was used for read trimming and porechop (0.2.4) (https://github.com/rrwick/Porechop) was used to remove adapter sequences (Bonenfant et al. 2022).

The long-read data was assembled with Flye (2.8.3) (Kolmogorov et al. 2020) run with the read input flag “--nano-raw” and the flag “--meta”. The resulting *de novo* assembly was polished using Illumina short read data from (Derelle et al. 2016) using Pilon (1.24) (Walker et al. 2014). Polishing was performed iteratively three times. The resulting assembly was run through the NCBI Foreign Contamination Screen (FCS) (Astashyn et al. 2024) prior to running downstream analysis. Ploidy level was assessed using Genomescope2 (2.0) (Ranallo-Benavidez et al. 2020).

#### Anvi’o metagenomics workflow and metabolic pathway analysis

The general Anvi’o (v7) workflow can be found here: https://merenlab.org/2016/06/22/anvio-tutorial-v2/. Briefly, the long-reads used for the initial assembly were aligned back to the final assembly using LongReadAligner (lra, 1.1.2) (Ren and Chaisson 2021). This allowed the calculation of read recruitment for future bins and for Anvi’o to perform hierarchical clustering. The resulting SAM file was then converted to a BAM file and a BAM index file using samtools (1.12) (Li et al. 2009). From there, Anvi’o (Eren et al. 2015) was used to create a contigs database (anvi-gen-contigs-database), followed by searching for Hidden Markov Models (HMMs) of rRNA genes and single-copy genes for protists and bacterial lineages (anvi-run-hmms). An Anvi’o profile (anvi-profile) was then created for the contig database to allow the use of the Anvi’o interactive interface for supervised binning (anvi-interactive (Eren et al. 2015)). Binning was performed manually based on read clustering, presence of rRNA genes, differences in GC content and read coverage. MAGs were extracted by summarising the binning effort (anvi-summarize).

Further analysis of the bacterial MAGs was performed with the Anvi’o suite of analysis programs. A contig database was created for each MAG (anvi-gen-contigs-database) before repeating the HMM search for rRNA genes and single-copy genes (anvi-run-hmms). The level of completeness was assessed for each bin by running anvi-estimate-genome-completeness, which measures the number of single-copy orthologues identified compared to the full set. Taxonomic assignment for each bin was performed with anvi-estimate-scg-taxonomy which uses single-copy core gene hits to the Genome Taxonomy DataBase (GTDB) (Parks et al. 2021) to assign taxonomy (with the exception of the Eukaryotic bin known to be *I. marina* and confirmed by blast hits to the 18S sequence on the NCBI).

Although Anvi’o performs *ab initio* gene calling on binned contigs, using the program prodigal (Hyatt et al. 2012), this is optimised for prokaryotic gene calling. Instead, to achieve an appropriate set of gene calls for the eukaryotic MAG, contigs were submitted to the Augustus web-server (https://bioinf.uni-greifswald.de/webaugustus/). Subsequent assessment of genome completeness was performed using BUSCO (v5 – stramenopile_odb10). Prediction of subcellular localisation of proteins was conducted using HECTAR (Gschloessl et al. 2008).

#### Searching Tara Oceans eukaryotic metagenomes (MAGs) and Single-Cell Assembled genomes (SAGs)

The DSYB protein sequence from *I. marina* was searched against the EUK_SMAGs database via the *Tara Ocean* Gene Atlas portal (Vernette et al. 2022), using a stringent e-value cut-off of 1e × 10^-70^, with abundance plotted as the percent of total reads. DSYB hits were further scrutinised by constructing maximum likelihood trees together with the *I. marina* query sequence as well as DSYB sequences of eukaryotic phytoplankton and bacterial taxa from (Curson et al. 2017; Curson et al. 2018), including functionally validated sequences. Geographic distributions were subsequently plotted for surface waters (SRF) and the deep chlorophyll maxima (DCM).

#### Maximum Likelihood tree construction of DSYB sequences

Multiple sequence alignments of DSYB/DsyB protein sequences (Curson et al. 2017; Curson et al. 2018) were generated using MAFFT (7.520) (Katoh et al. 2002). Alignments were trimmed in auto mode by trimal (1.4.1) (Capella-Gutiérrez et al. 2009), where sites made up of 60% gaps were removed. A maximum likelihood tree was then produced with IQtree (Minh et al. 2020) with the following parameters: iqtree -s infile.txt -bb 10000 -safe -bnni -alrt 10000 -st AA - seed 1000 -msub nuclear -t RANDOM -nt AUTO -pre out -m TEST (Q.pfam+I+R5 was determined to be the best fit model). The raw FASTA file of all sequences used to construct the phylogenetic tree is given in **Dataset S1**. The final trimmed alignment used for tree construction was 365 amino acids in length.

#### Genome annotations

Genome annotation was used to compare the functional landscape of *I. marina* against other more widely studied stramenopiles. The genomes of *Phaeodactylum tricornutum*, *Schizochytrium aggregatum* ATCC 28209*, Phytophthora sojae, Cafeteria roenbergensis* BVI *and Blastocystis hominis* (Singapore Isolate B) were acquired from the JGI genome browser. The eggNOG-mapper annotation server (Cantalapiedra et al. 2021) was used to run gene annotation on each genome using default settings. Subsequent KEGG Orthology annotation data was plotted using the venny4py python package (https://github.com/timyerg/venny4py).

Estimation of the metabolic capacity of organisms in the community, both eukaryotic and prokaryotic, was achieved using the KEGG orthologue annotation from ghostKOALA (Kanehisa et al. 2015) and the KEGG Mapper-Reconstruct tool (https://www.genome.jp/kegg/mapper/reconstruct.html) to overlay KEGG orthologues on biosynthetic pathways, followed by manual inspection to determine whether they were likely complete. Taxonomic assignment of the closest related KEGG orthologue for each annotated protein was also extracted from ghostKOALA output and used to identify protein sequences with close similarity to red algal protein sequences.

## Results and Discussion

### Imaging and physiology of the *I. marina*-bacterial consortium

The Nanomonadea species *I. marina* (CCAP 977/1) was originally isolated from an estuarine environment with a consortium of bacteria (Cavalier-Smith and Scoble 2013), with which it has been maintained ever since. Due to the lack of cultured MASTs, microscopy studies of these organisms are limited. We therefore first inspected I. *marina* with its associated bacterial assemblage using bright-field microscopy. A mixed population of protistan and bacterial cells was clearly visible (**Figure S1A**). Scanning electron microscopy (SEM) revealed physical associations between *I. marina* and bacteria, including bacterial attachment along the length of the protistan flagellum (**Figure 1A, left image**). We also observed clumping and biofilm formation, particularly in older cultures (**Figure S1B-D**), as well as evidence of dead and decaying *I. marina* cells, with associated bacteria in close proximity, potentially feeding on the *I. marina* biomass (red arrow, **Figures S1E**). Our images corroborate earlier descriptions of *I. marina* as a nanoflagellate of 2-3 µm with a single smooth flagellum (Cavalier-Smith and Scoble 2013) (**Figure 1A; Figure S1F**). Additionally, we identified the presence of an opening at the flagellum base where it joins the cell body (red arrow) (**Figure 1A, right image**). Initial analysis of the bacterial community by plating on marine broth agar identified 4 morphologically different colonies. Analysis of the 16S rRNA genes from samples of these colonies identified them as *Ruegaria sp.*, *Gammaproteobacteria bacterium*, *Winogradskyella sp.*, *Marinobacter hydrocarbonoclaticus* and *Pseudooceanicola marinus*.

**Figure 1.**
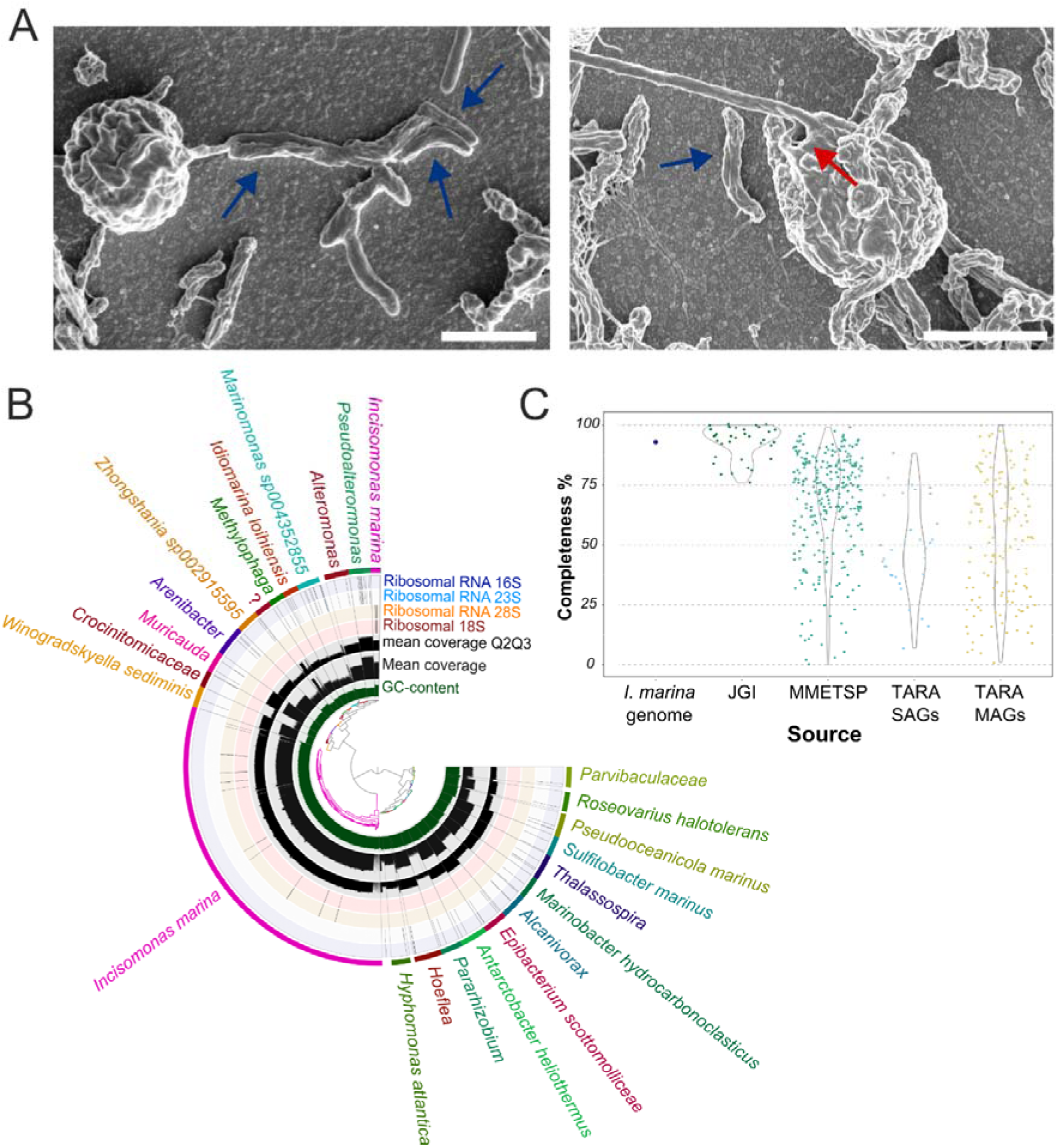
Imaging and metagenomic analysis of *Incisomonas marina* and its associated bacterial community. **A.** Scanning electron microscopy of *I. marina*. Individual *I. marina* cells displaying a single flagellum as well as physically close associations with bacteria, including along the length of the flagellum; blue arrows indicate bacterial cells. Red arrows indicate a consistently observed opening at the base of the flagellum, as well as an ‘overhanging lip’ as observed by [13]. Scale bar: 2 µm. **B**. A view of the Anvi’o interactive interface used to perform supervised binning for metagenome assembled genome (MAG) recovery. The central tree is a hierarchical clustering of re-aligned long reads to the final metagenome assembly. Each track surrounding this tree represents (in order from inside out): GC content, mean coverage (read recruitment), mean coverage for Q2 and Q3, locations of rRNA genes for 18S, 28S, 23S and 16S. Each bin is coloured individually by the final outermost track with an associated taxonomic prediction where possible. **C**. Comparison of the genome completeness estimations for available stramenopile genomes and transcriptomes (the latter from the Marine Microbial Eukaryote Transcriptome Project, MMETSP [49] versus the estimated completeness of the *I. marina* MAG generated in this study (93%). Measured by BUSCO (v5) [51] against stramenopile_odb10.

To investigate the dependency of *I. marina* on the bacteria present versus the organic material provided from the culture medium (ASWP medium that includes soil extract as well as a boiled barley grain), cultures grown in ASWP were treated with two different antibiotic cocktails (**Table S1**). The cultures were imaged and compared to a control with no antibiotics, as well as another media-only control (no cells), over a time course of 7 days. After 5 days of antibiotic treatment no *I. marina* cells were visible (**Figure S2A-B**). Likewise, no bacterial growth was observed on marine broth agar plates inoculated with antibiotic treated cultures, compared to the untreated control (**Figure S2C**). Whilst it is possible that the antibiotics killed *I. marina* directly, the antibiotics chosen for this study have been used widely to minimise bacterial contamination of other stramenopile cultures (Droop 1967; Guillard 2005; Wilkens and Maas 2012; Koedooder et al. 2019; Ferrante et al. 2020) and we used lower concentrations than those reported in the literature (**Table S1**). Thus, these observations suggest that removal of the bacterial community from the culture by antibiotics resulted in loss of a nutritional source from *I. marina*, leading to death of the protist.

### Metagenomics analysis of *I. marina* and its associated bacterial community

To investigate further *I. marina* and its microbiome, we carried out metagenomic sequencing of the entire consortium, using Oxford Nanopore long-read sequencing. This yielded 1,415,560 reads spanning a total of 16,635,245,180 bases. A supervised binning effort performed in conjunction with the Anvi’o taxonomic assignment tool (Eren et al. 2015) generated 24 bins, accounting for 97.16% of all nucleotides (**Table S2-S3**). All but two of the bins were assigned as being of bacterial origin, confirmed by the presence of 16S/23S ribosomal RNA sequences and the single-copy core gene hits to the GTDB (Parks et al. 2021) (**Figure 1B; Table S4**). It was possible to recover 22 bacterial genomes as near complete MAGs, ranging in size from 2.9 Mbp to 5.5 Mbp (**Table S3**); a further bin was incomplete and not taxonomically assigned (**Table S4**). Of the 22 complete/near complete bacterial MAGs, ten were *Alphaproteobacteria* (including five *Rhodobacteraceae*) and eight *Gammaproteobacteria* (including three *Alteromonadaceae* species) (**Table S4**). The remaining four genomes were from *Bacteroidia* (all *Flavobacteriales*). Notably, the described bacterial community included many taxa commonly found associated with photosynthetic stramenopile relatives, the diatoms (namely *Rhodobacteraceae*, *Alteromonadaceae* and *Flavobacteriales*) (Amin et al. 2012). Eight of the MAGs (of the genera *Pseudoalteromonas, Alteromonas, Methylophaga, Arenibacter, Muricauda, Alcanivorax, Pararhizobium, Hoeflea and Thalassospira*) had no species level taxonomic prediction suggesting potentially novel species, and two bins had taxonomic prediction only to family level (*Crocinitomicaceae* and *Parvibaculaceae*). As an indication of the quality of our metagenomic assembly for these bacteria, 18 of the genomes were >97% complete, and seven comprised a single contig (**Table S3**).

The remaining two bins both had 18S/28S rRNA genes that were identified as *I. marina*, alongside the same GC content, and so they were combined to form the *I. marina* MAG (hereinafter referred to as genome) (**Table S3 and S4**). The *I. marina* genome was determined to be haploid by calculating k-mer frequencies from *I. marina* specific reads (**Figure S3**). A phylogenomic tree showing the relationship of *I. marina* to other stramenopiles is given in **Figure S4**. With a genome of ∼68 Mbp (**Table S3**), the *I. marina* genome is similar in size to many other sequenced stramenopile genomes including *Schizochytrium* (now *Aurantiochytrium*) species (Liang et al. 2020) and several *Phytophthora* species (Tyler et al. 2006; Lamour et al. 2012; Liang et al. 2020), but larger than the diploid diatoms *Thalassiosira pseudonana* (32.4 Mbp; (Armbrust et al. 2004)) and *Phaeodactylum tricornutum* (27.4 Mbp; (Bowler et al. 2008) and other heterotrophic stramenopiles *Cafeteria roenbergensis* BVI (36.3 Mb; (Hackl et al. 2020)) or *Blastocystis hominis* Singapore Isolate B (18.8 Mb; Denoeud et al., (2011). In fact, genome size varies considerably even within individual stramenopile groups, with an almost 50-fold difference amongst diatoms, from 33 Mb for *Cyclotella nana* to 1.5 Gb in *Thalassiosira tumida* (Roberts et al. 2024) and even ranging 32 Mb - 295 Mb for the *Phytophthora* genus (McGowan and Fitzpatrick 2020). These variations are likely due mainly to a higher proportion of repetitive sequences in the larger genomes. In this context, analysis of the repetitive sequence content of the *I. marina* genome using Repeatmasker (Tarailo-Graovac and Chen 2009) determined that 20.5% was made up of repeats. This consisted predominantly of simple repeats (5,834,774 bp – 8.57%) and retroelements (2,698,197bp - 3.96%, including 1.27% large tandem repeat elements), as well as DNA transposons (1.23%).

To investigate the gene complement of *I. marina,* the pipeline shown in **Figure S5** was followed, with the gene-calling software Augustus (Stanke and Morgenstern 2005) predicting a total of 20,091 gene models. BUSCO analysis subsequently determined the genome completeness of this predicted gene set to be 93%. A meta-comparison with other sequenced stramenopiles (including 498 accessions from *Tara* (Delmont et al. 2022), Marine Microbial Eukaryote Transcriptome Sequencing Project (MMETSP) (Keeling et al. 2014) and Joint Genome Institute) places the *I. marina* genome in the top 10% for genome completeness using BUSCO (Manni et al. 2021) (**Figure 1C**). It should be noted that this analysis compares different types of ‘omics dataset (e.g. genomes of cultured strains versus transcriptomes), but nevertheless gives an indication of the quality of the *I. marina* genome retrieved. Gene annotation analyses revealed the functional repertoire of *I. marina* is comparable to other sequenced stramenopiles (**Figure S6**). Notably the most frequent annotation is “Function Unknown”, and many genes have no annotation at all, so only 20% of *I. marina* genes have a known function. Nevertheless, 1545 genes were annotated with a COG annotation, with post-translational modification (O), translation (J), carbohydrate (G) and amino acid transport/metabolism (E) being the most well-represented categories in both heterotrophic and photosynthetic stramenopiles (**Figure S6**). Comparative genomics of shared and unique KEGG orthologs (KOs) between a diverse selection of stramenopiles revealed that the largest set of KOs was shared between each species, likely representing core metabolic functions. However, *I. marina* had the greatest number of unique KOs (504) of the six represented species, which suggests that this species possesses a unique metabolism within the sampled stramenopiles (**Figure 2**). Amongst these unique proteins, included enrichment of 19 lysosome-associated proteins, including four cathepsin proteases, as well as 3 branched-chain amino acid transporters.

**Figure 2.**
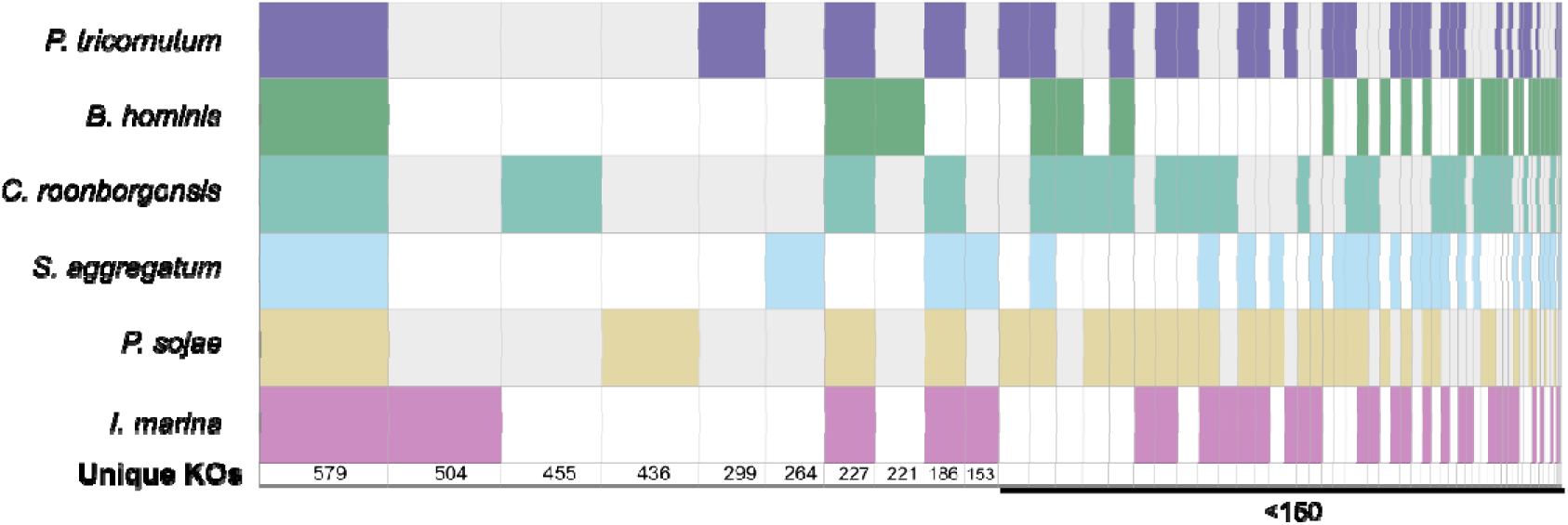
Shared and unique KEGG Orthologues of *Incisomonas marina* with other stramenopiles. SuperVenn diagram of shared and unique predicted proteins (by KEGG orthology) of *I. marina* with *Phaeodactylum tricornutum*, *Phytophthora sojae*, *Schizochytrium aggregatum, Cafeteria roenbergensis* and *Blastocystis hominis*. These species represent groups across the stramenopiles: Nanamonadea, Bacillariophyceae, Oomycota, Labyrinthula and bikosia respectively. Note that sets with below 50 values are not shown.

The presence of a chloroplast of red-algal origin in diatoms, which is also shared with many other algal groups including dinoflagellates and haptophytes, led to the proposal of a single secondary endosymbiosis, the so-called Chromalveoate Hypothesis (Cavalier-Smith 1999). However, phylogenomic analysis of nuclear genomes from these different algal groups does not resolve them as monophyletic (Strassert et al. 2019) and instead it is likely that multiple secondary and tertiary endosymbioses have occurred over evolutionary history (Dorrell and Smith 2011; Sibbald and Archibald 2020). Indeed, in searching for evidence of a red-algal like plastid in the genome of *I. marina*, of 4208 “Protist” annotated genes only 74 had unique red algal annotated gene calls and only one was identified as having a predicted protein product with localisation to the chloroplast of photosynthetic species **(Table S5)**. This analysis suggests that it is unlikely that the basal MAST-3 lineage of stramenopiles had chloroplasts at any point during their evolutionary history.

### Examining the requirement of *I. marina* for essential vitamins and amino acids

To advance knowledge of the nutritional requirements of *I. marina*, we took a more targeted approach. Given the widespread vitamin auxotrophy amongst algal lineages (Croft et al. 2006; Helliwell 2017) and poor understanding of protist vitamin requirements (Needham et al. 2022), we examined the *I. marina* genome for genes encoding biosynthetic enzymes for B vitamins, which provide essential enzyme cofactors. *I. marina* was predicted to encode complete biosynthesis pathways for B_3_ (niacin), B_5_ (pantothenate) and B_9_ (folate) (**Figure 3**, left-hand column) and be auxotrophic for B_1_ (thiamine), B_2_ (riboflavin), B_6_ (pyridoxine) and B_7_ (biotin). The presence of the B_12_-dependent isoform of methionine synthase (*METH*), but not the B_12_-independent isoform (*METE*) (Helliwell et al. 2011), suggests that *I. marina* also requires an exogenous supply of B_12_ (cobalamin). Additionally, we identified that *I. marina* encodes both L-galactonolactone dehydrogenase (GLDH) and L-gulonolactone oxidase (GULO) for vitamin C (ascorbate) biosynthesis (**Table S6**). This was also the case for C. *roenbergensis,* but not *S. aggregatum*, *P. sojae*, *P. tricornutum* and *B. hominis*. In plants and algae, GLDH has functionally replaced GULO, likely due to its ability to decouple ascorbate biosynthesis from H_2_O_2_ production, promoting the role of ascorbate as a photoprotective antioxidant (Wheeler et al., 2015). The selective pressure to retain vitamin C biosynthesis in *I. marina* may have been driven by the inability of bacteria to synthesise this essential cofactor and antioxidant, which could also mitigate oxidative stress resulting from biotic interactions (Sunagawa et al., 2009). Thus, whilst *I. marina* clearly requires certain vitamins, contrary to prevailing views on the nutritional dependencies of heterotrophic protists, *I. marina* has the genes necessary to synthesise several vitamins itself and thus will likely contribute to the cycling of such crucial currencies in marine microbial communities.

**Figure 3.**
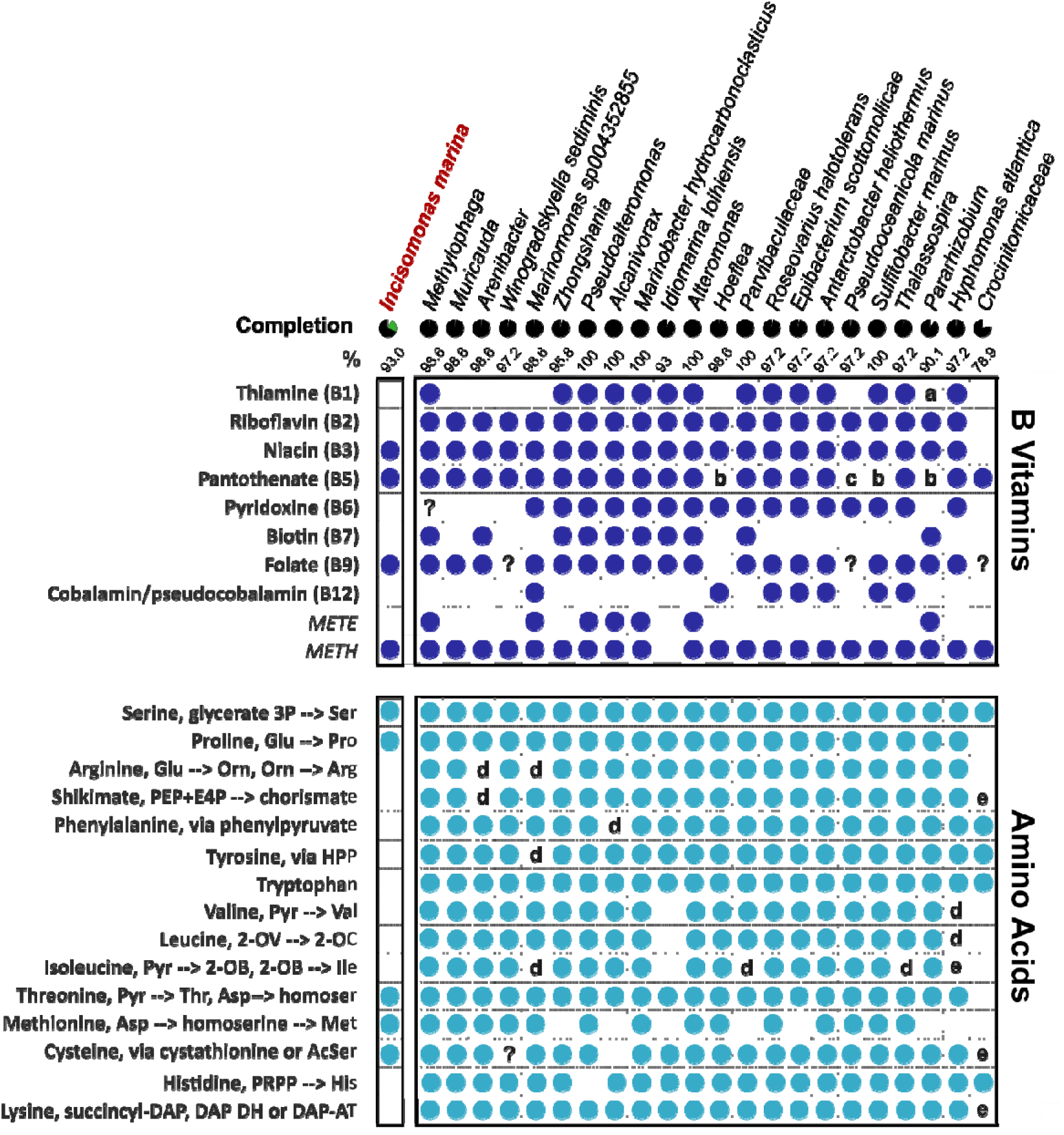
*Incisomonas marina* lacks *de novo* biosynthesis pathways for multiple B vitamins and amino acids. Predictions of B vitamin (**top**) and amino acid (**bottom**) biosynthesis pathways in each metagenome assembled genome where a circle represents instances where a pathway is predicted to be complete (or the gene to be present in the case of B_12­_-independent and dependent methionine synthases (*METE*/*METH,* respectively)). ‘a’ indicates where the enzyme ThiL is not present, ‘b’ where PanB and PanC are missing and ‘c’ where only PanB is missing. For the amino acid biosynthesis analysis, ‘d’ and ‘e’ indicates where a partial biosynthesis pathway was identified but with 1 or 2 enzymes of that pathway missing respectively, as predicted by KEGG. Ambiguous biosynthesis capability is labelled ‘?’.

*I. marina* encodes the genes necessary for processing ammonia into organic nitrogen i.e. via glutamine synthase and glutamine oxoglutarate aminotransferase (GOGAT) (**Table 1**). We also found genes for enzymes of the urea cycle, previously identified in diatoms (Allen et al. 2011), and thought to play a role in adaptation to limiting nitrogen conditions. In *P. tricornutum* and *T. pseudonana*, the enzyme that catalyses the first step, carbamoyl phosphate synthase (CPS), is located in the mitochondria (Allen et al. 2011). Diatoms also encode a second CPS that lacks an organelle targeting sequence and is presumed to be involved in cytosolic pyrimidine biosynthesis. In a similar manner, we identified two hits for CPS in *I. marina*, one of which is predicted to be mitochondrially-targeted (**Table 1**). Reciprocal BLAST of these sequences against the *T. pseudonana* genome yielded both diatom CPS genes (JGI protein ids: 24248 and 24195). *I. marina* also encodes an arginase, necessary for production of urea from arginine, as well as ornithine and proline. However, it lacks the capability to synthesise *de novo* many of the protein amino acids (**Figure 3**). Whilst there were predicted genes for all enzymes of proline, methionine and cysteine biosynthesis, as well as those derived directly from intermediary metabolism (glycine, alanine, glutamate, glutamine, aspartate, asparagine, serine and threonine), those of pathways for lysine, arginine, the aromatic amino acids (phenylalanine, tyrosine, tryptophan) or the branched-chain amino acids (valine, leucine, isoleucine) were either entirely missing or incomplete. The absence of the shikimate pathway enzymes for the synthesis of chorismate is notable given that *I. marina* encodes the subsequent enzymes to produce folate (B_9_) from chorismate.

**Table 1.** N assimilation and urea cycle enzymes identified in the genome of *Incisomonas marina*.

| Process | Enzyme | EC No. | KEGG orthologue No. | *Incisomonas marina* protein Id | Predicted targeting category |
| --- | --- | --- | --- | --- | --- |
| N assimilation | Glutamine synthetase | 6.3.1.2 | KO1915 | g18223.t1 | Other |
|  |  |  |  | g7363.t1 | Mitochondrial |
|  |  |  |  | g14366.t1 | Other |
|  | Glutamine oxoglutarate aminotransferase (GOGAT) | 1.4.1.13 | KO0265 | g9516.t1 | Mitochondrial |
| Urea Cycle | Arginase | 3.5.3.1 | KO2965 | g12696.t1 | Other |
|  | Carbamoyl phosphate synthase (CPS) | 6.3.4.16 | KO1948 | g18404.t1 | Mitochondrial |
|  |  | 6.3.5.5 | KO0370 | g8254.t1 | Other |

The high level of completeness of the genome (93%) suggests the lack of homologues of genes for both vitamin and amino acid biosynthesis is likely to be correct, although it does not rule out divergent forms of the enzymes or indeed alternative biosynthetic routes (Webb et al. 2007), which would not be detected in this KEGG orthologue analysis. Nonetheless, it can be concluded that *I. marina* needs to obtain a suite of essential metabolites for normal growth. Identification of a complete urea cycle as well as branched chain amino acid transporters, would enable *I. marina* to acquire and detoxify organic compounds obtained through digestion of bacteria.

To assess the potential for the associated bacterial consortium to provide *I. marina* with its required amino acids and B-vitamins, we investigated the metabolic capabilities of the sequenced members. With regards the B-vitamins, just three species (*Pseudoalteromonas* sp., *Alcanivorax* sp., *and Marinobacter hydrocarbonoclasticus*) were autotrophic, since they were predicted to produce B_1_-B_9_ and encoded the B_12_-independent METE (**Figure 3**, right-hand columns). In contrast, all the other bacteria appeared to have a requirement for an external source of at least one vitamin. *Crocinitomicaceae* was predicted to produce just one B-vitamin, pantothenate (B_5_), although its genome was less than 80% complete. Of species with >90% completion, *Pararhizobium* sp. and *Hoeflea* had just four predicted biosynthesis pathways. Over half the bacteria were missing all four required enzymes for biotin (B_7_) biosynthesis, whereas for those species unable to synthesise pantothenate (B_5_), one lacked just the first committed enzyme, PanB (*Pseudooceanicola*) and three others (*Hoeflea*, *Pararhizobium* and *Sulfitobacter*) lacked both PanB and the last enzyme, PanC.

By comparison, the majority of bacteria encoded the genes necessary for biosynthesis of the protein amino acids (**Figure 3**, right-hand columns). As expected, all species synthesising lysine used the typical eubacterial succinyl-DAP route, rather than the DAP aminotransferase characteristic of eukaryotes, and whilst the majority used the acetylserine route for cysteine biosynthesis, *Muricauda* and *Arenibacter* encoded enzymes for the alternative route via cystathionine. *Crocinitomicaceae* lacked most enzymes for biosynthesis of proline, arginine, methionine and branched-chain amino acids as well as the shikimate pathway, but again this might be due to low genome completeness. In contrast, the absence of genes for branched-chain amino acid biosynthesis in *Hyphomonas atlantica* and *Idiomarina loihiensis* is likely to be the case, since their genomes were 97% & 93% complete, respectively. The capacity for methionine biosynthesis was observed the least frequently amongst the bacteria, whereas *I. marina* can do so. Together, our analysis provides evidence that *I. marina* can satisfy its requirements for several essential organic nutrients from its co-habiting bacteria, and that cross-feeding may also occur between bacterial community members.

### Identification of an *I. marina DSYB* gene for biosynthesis of potent bacterial chemoattractant DMSP

DMSP is an abundant marine organosulfur compound with important roles in stress protection and signalling (Kuhlisch et al. 2023). Produced by many bacteria (Curson et al. 2017) and eukaryotic phytoplankton (Curson et al. 2018), DMSP gives rise to the climate-active gas, dimethylsulphide (DMS). DMSP is typically produced through the transamination pathway (Curson et al. 2017), via *S*-adenosylmethionine (SAM)-dependent 4-methylthio-2-hydroxybutyrate (MTHB) *S*-methyltransferase, encoded by the DSYB/DsyB gene (Curson et al. 2017; Curson et al. 2018) (**Figure 4A**), although an alternative algal biosynthesis enzyme, DSYE, has recently been identified (Wang et al. 2024).

**Figure 4.**
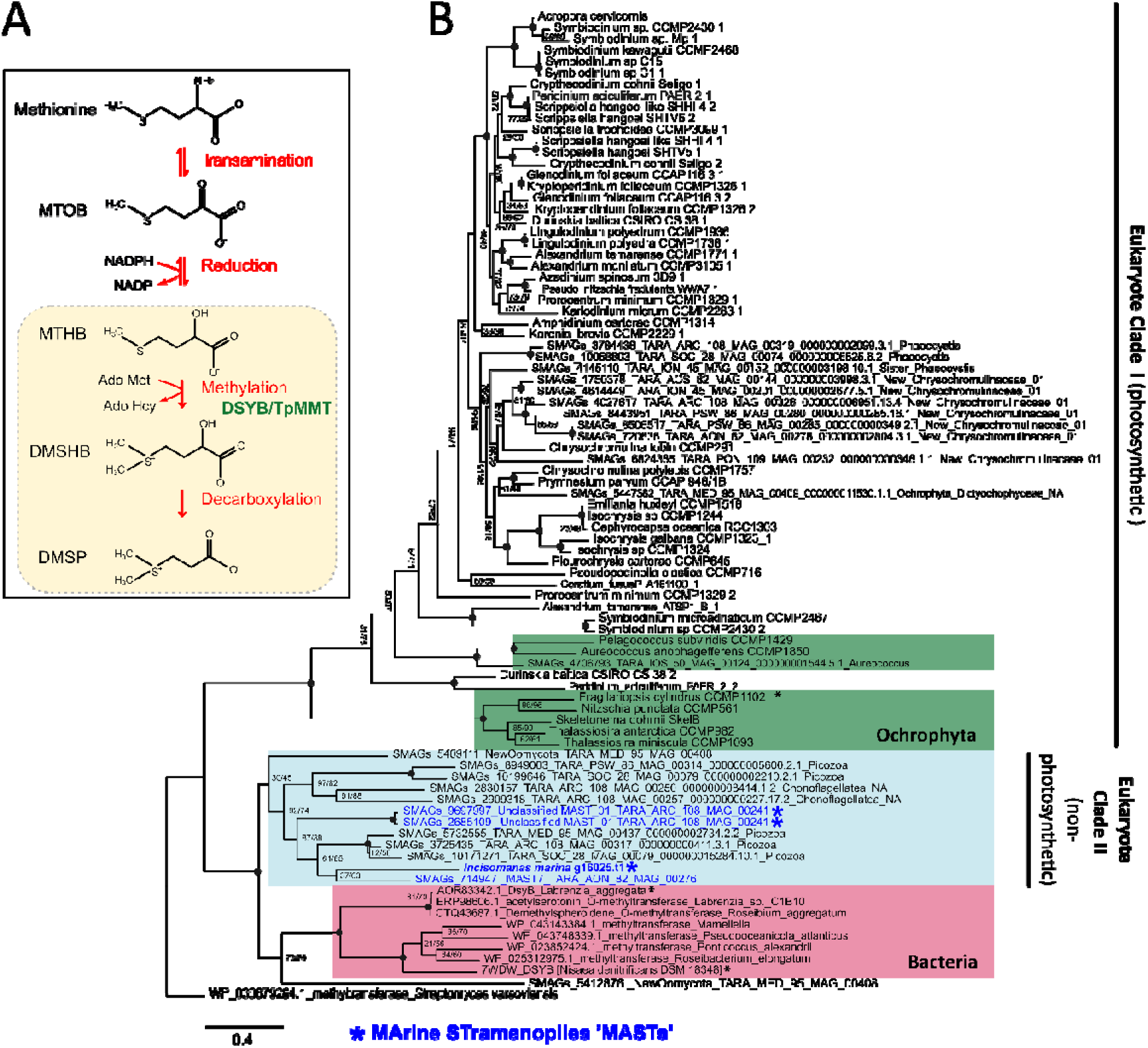
Identification of DMSP biosynthesis enzyme DSYB in *Incisomonas marina* and several other MASTs. **A.** Schematic diagram of the transamination pathway for DMSP biosynthesis from methionine. The key biosynthesis enzyme S-adenosylmethionine (SAM)-dependent 4-methylthio-2-hydroxybutyrate (MTHB) S-methyltransferase DSYB, is indicated in green. DSYB has been identified in heterotrophic bacteria as well as certain eukaryotic algae [30, 31, 63]. The MMT enzyme identified by Kagyama et al., (2018) in *Thalassiosira pseudonana* is an alternative proposed methylation enzyme. MTOB – 4-methylthio-2-oxobutyrate; MTHB – 4-methylthio-2-hydroxybuterate; DMSHB – 4 dimethylsulphonio-2-hydrxybutyrate. Pathway constructed from [31, 64]. In eukaryotic algae DSYB enzyme and DMSP synthesis is localised to both chloroplasts and mitochondria (yellow box) [31]. **B**. Maximum likelihood gene tree of DSYB protein sequences from bacteria, eukaryotic phytoplankton, and new hits identified in this study for *I. marina* and other MASTs (blue). MAST sequences form a new clade of eukaryote DSYB proteins (‘Eukaryote Clade II’) together with hits from several other heterotrophic eukaryotes (from Picozoa, New Oomycota and choanoflagellates) retrieved from the *Tara Oceans* eukaryotic metagenome and single-cell assembled genome databases. The new Eukaryote Clade II is sister to the bacterial clade of DSYBs and sits apart from Eukaryote Clade I that contains eukaryotic phytoplankton DSYB sequences from [31]. Three DSYB enzymes that have been functionally validated are labelled with an ‘*’. The non-DSYB methyltransferase sequence of *Streptomyces varsoviensis* (WP_030879264.1) was used as an outgroup. Groups of related organisms are indicated with coloured boxes. IQtree was used in model prediction mode to create a maximum-likelihood tree using ultrafast bootstrapping and the -altr flag, resulting in two values of node support. Q.pfam+I+R5 was determined to be the best fit model. The final trimmed alignment used for tree construction was 365 amino acids in length. Node support values were removed above 90/90 are indicated with a filled black circle.

In photosynthetic algae, DSYB is localised to chloroplasts and mitochondria (Curson et al. 2018). Previous searches of heterotrophic stramenopiles (including Labyrinthulea *Aplanochytrium* (sp. PBS07 and *A. stocchinoi* GSBS06), *Aurantiochytrium limacinum* ATCCMYA1381 and *S. aggregatum* ATCC28209) did not yield hits for *DSYB* (Curson et al. 2018). Nor did we find the gene in *B. hominis* Singapore isolate B (sub-type 7) or *C. roenbergensis* BVI. However, querying the *I. marina* genome with a diatom sequence (*Fragilariopsis cylindrus*, protein 238045) (Curson et al. 2018), yielded a robust hit (that did not resemble *F. cylindrus* DSYE (Wang et al. 2024)). Moreover, a mitochondrial transit peptide was identified. To examine the presence of putative DSYB sequences in other MASTs, we used the *I. marina* sequence to query the *Tara Oceans* eukaryotic metagenome and SAG database (EUK_SMAG) (Vernette et al. 2022). We retrieved three robust hits for MASTs (all hits with an e-value cut-off below 1e × 10^-70^ are shown in **Table S7**). A further two were obtained from a MAG taxonomically assigned as ‘New Oomycota’ (i.e. heterotrophic stramenopile taxa) (Delmont et al. 2022). Our search also yielded sequences from choanoflagellates and Picozoa, as well as eukaryotic phytoplankton, with many predicted to be mitochondrially localised (**Table S7**). A further two hits for heterotrophic Picozoa were obtained by mining additional SAG databases (Schön et al. 2021) (F_COSAG_04_NODE_10F_COSAG02 and F_COSAG04_NODE_1660). Construction of a maximum likelihood tree revealed that the MAST sequences (labelled blue) grouped together with robust support (**Figure 4B**), along with sequences from eukaryote taxa; in all cases predicted to be heterotrophic (**Table S7**) (Delmont et al. 2022). This new clade of heterotrophic eukaryotic DSYBs (‘Eukaryote Clade II’, **Figure 4B**) groups separately from the established eukaryotic clade (‘Clade I’) containing algal DSYBs (Curson et al. 2018), and is instead sister to the bacterial clade (Curson et al. 2017; Li et al. 2022). A multiple sequence alignment of functionally verified DSYB/DsyB enzymes, demonstrates that the MAST sequences have key residues experimentally determined to be necessary for DSYB/DsyB function, including for MTHB and SAM binding (**Figure 5**), indicating that they are likely functional.

**Figure 5.**
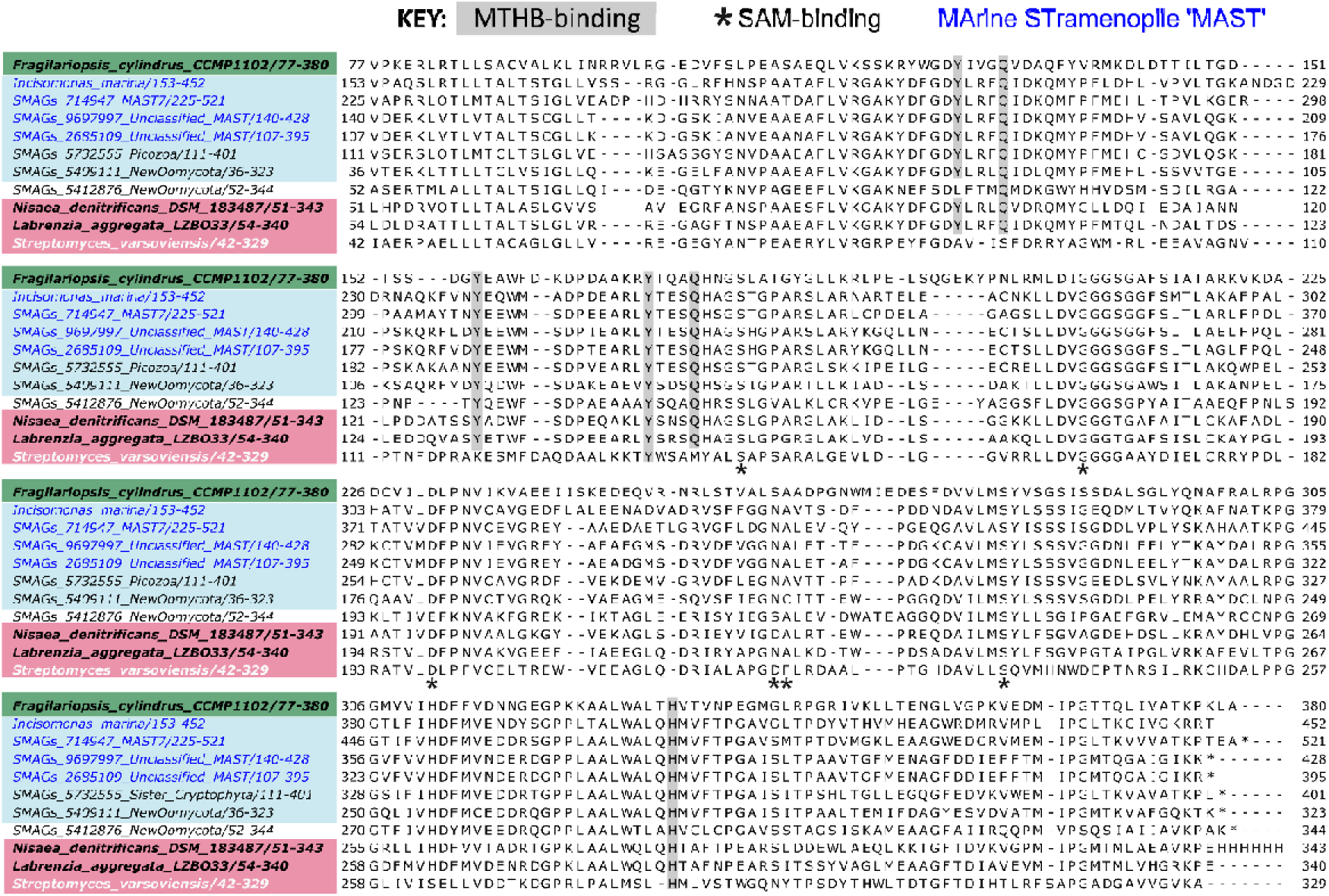
Multiple sequence alignment of bacterial and eukaryotic DsyB/DSBY protein sequences. DsyB protein sequences that have been cloned and functionally characterised from the bacteria *Nisea denitrificans* (NCBI identifier: 07WDW_A) [64], *Labrenzia aggregata LZB033* (NCBI identifier: AOR83342) [32] and the diatom *Fragilariopsis cylindrus* (JGI protein ID: 238045) [31] are highlighted in bold. The sequence of a ‘non-DsyB’ methyl-transferase enzyme from *Streptomyces varsoviensis* (NCBI identifier: WP_030879264.1) incapable of DMSP biosynthesis is white. Sequences from *Incisomonas marina* and representatives from ‘Eukaryote Clade II’ shown in Figure 4B are also included, as well as both ‘New Oomycota’ sequences. Taxa are coloured according to their phylogenetic grouping from the maximum likelihood tree in Figure 4B. Residues involved in MTHB binding and SAM binding are labelled as indicated.

Plotting the distribution of the MAST SMAG hits revealed a cosmopolitan distribution in marine systems globally of the DSYB MAST-7 hit (**Figure 6A-B**), whereas the ‘Unclassified MAST’ sequences showed a more confined biogeography (**Figure 6C-D**). Nevertheless, the identification of convincing DSYB sequences in MAST lineages as well as from other heterotrophic eukaryotes suggests the contribution of such protists to global DMSP cycling is likely more significant than previously recognized. Given that DMSP is a potent chemoattractant and important nutrient source for many marine bacteria (Seymour et al. 2010), DMSP biosynthesis could thus enable MASTS to attract and sustain the bacterial community.

**Figure 6.**
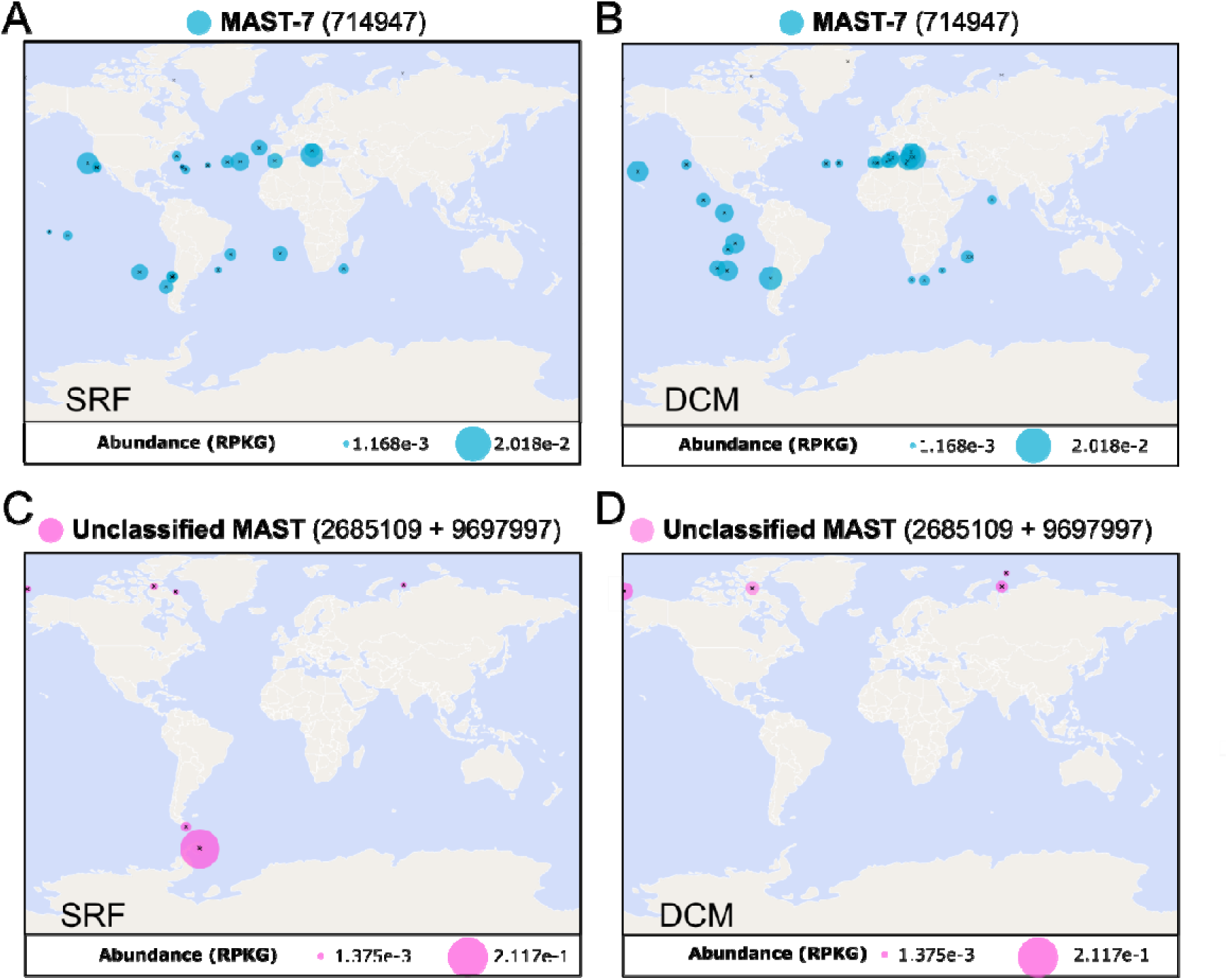
Global abundance of MAST DSYB in marine systems. Bubble plots showing abundance (as a percentage of total reads) of ‘MAST-7 hit’ SMAG_714947 in surface waters (SRF) (**A**) and the deep chlorophyll maxima (DCM) (**B**). Equivalent plots are also shown for ‘unclassified MAST’ hits (SMAG_2685109 and SMAG_9697997) (**C** and **D**). Data were compiled through querying Eukaryotic metagenomes (MAGs) and single cell assembled genomes (SAGs) via the Ocean Gene Atlas portal [30].

## Conclusion

Across the eukaryotic tree of life there is considerable bias in genomic sequencing (del Campo et al. 2014; Torruella et al. 2025). This is acutely apparent in the stramenopile group, which displays an enormous range of habitat, trophic and cell structural diversity and yet sequencing efforts focus mainly on photosynthetic lineages and terrestrial oomycetes (Armbrust et al. 2004; Bowler et al. 2008; del Campo et al. 2014; Kronmiller et al. 2023). In contrast, the basal lineages are poorly understood and understudied. This is despite the fact that MASTs are likely major players in ocean carbon cycling and the marine microbial loop (Massana et al. 2006) with recent work identifying MAST-3s as grazers of *Prochlorococcus,* one of the most globally abundant phytoplankton taxa (Wilken et al. 2023). Our study provides important evidence that nutritional demands for certain B vitamins and amino acids may well be key factors driving such predator-prey interactions. However, *I. marina* encodes genes necessary to synthesise several vitamins itself (vitamins C, B_3_, B_5_ and B_9_). Notably, our metabolic pathway analysis also reveals that no one member of the *I. marina* consortium is capable of synthesising all the organic nutrients examined. Metagenomic analysis of the ‘meta-metabolome’ thus suggests cross-exchange of nutrients is possible between members of this community, which has been maintained stably since its isolation from environmental samples. Further, our work reveals unexpected metabolic attributes of MASTs. Whilst not previously found in other heterotrophic stramenopile genomes, we identified the DMSP biosynthesis gene *DSYB* in *I. marina* as well as environmental MAST SMAGs. Our findings raise important questions regarding the evolution of DMSP biosynthesis in eukaryotes, and highlights a role for heterotrophic stramenopiles in global DMSP production and sulphur cycling. Finally, our study indicates that key metabolic innovations of ancestral heterotrophic stramenopiles (urea cycle, GLDH pathway for vitamin C biosynthesis, and DMSP biosynthesis) may have provided important foundations for the evolution of some of the most successful groups of phototrophic organisms on the planet.

## Acknowledgements

We acknowledge support from the University of Cambridge NERC DTP for a studentship for D.E.A. and a NERC Independent Research Fellowship grant NE/R015449/2 to K.E.H.

## Competing Interests Statement

The authors declare no competing interests.

## Data Availability

All metagenomics data for this study have been deposited to the European Nucleotide Archive (ENA) using the accession number: PRJEB86075.

## Notes

https://www.ebi.ac.uk/ena/browser/view/PRJEB86075

### Competing Interest Statement

The authors have declared no competing interest.

