## Supplementary Information for "Metagenomics of the MAST-3 stramenopile, *Incisomonas,* and its associated microbiome reveals unexpected metabolic attributes and extensive nutrient dependencies"

### Supplementary Figures and Tables

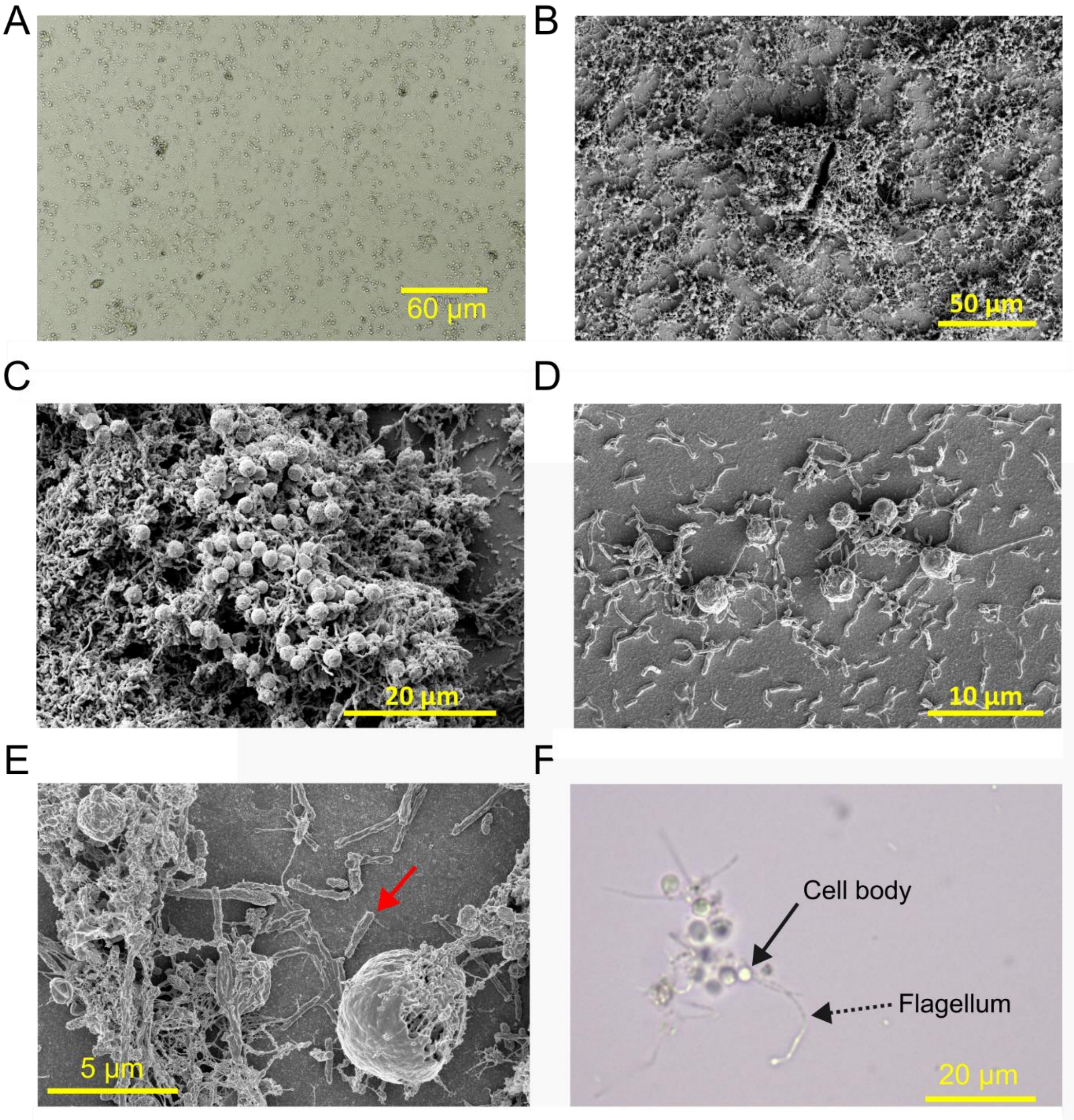


**Figure S1. Imaging of the *Incisomonas marina* CCAP 977/1 culture. A.** Light microscopy image of *I. marina* and bacteria (40× magnification). **B-D**. Scanning Electron Microscopy (SEM) images of *I. marina* and associated bacteria displaying benthic growth and clumping behaviour. **E**. SEM image of dead and decaying *I. marina* cells (e.g., bottom right quadrant) with associated bacteria in close proximity (red arrow). **F.** Oil immersion 100× magnification light microscopy image of *I. marina* showing typical clumps that form in the medium. The cell body and flagellum *of I. marina* are annotated with arrows.

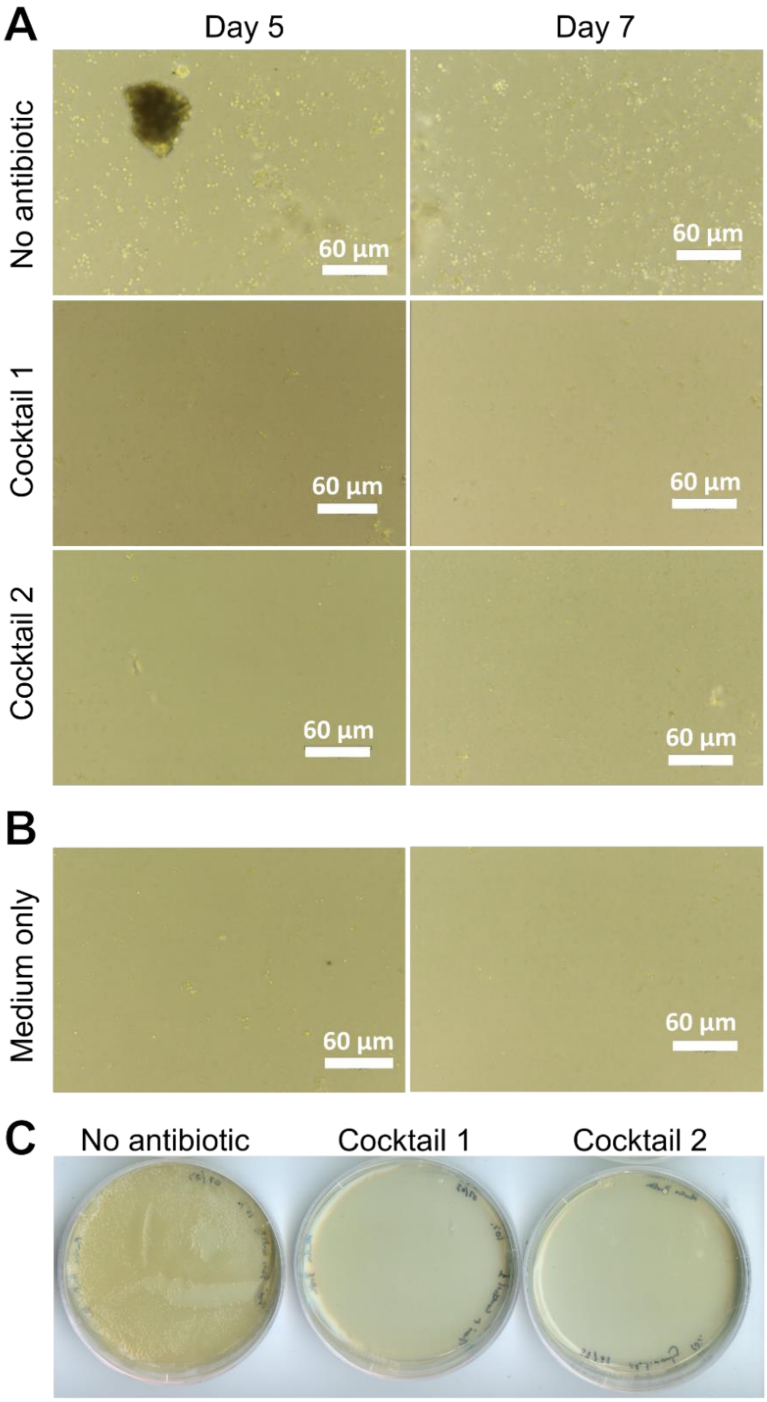


**Figure S2. Treatment of *Incisomonas marina* cultures with antibiotics eliminated growth of *I. marina* and bacteria. A.** Light microscopy images of *I. marina* cultures after 5- and 7-days treatment with two different antibiotic cocktails (**Table S1**), compared to the no antibiotic control. **B.** Images taken of 'media only' controls containing artificial seawater for protozoa (ASWP) medium + grain i.e., without inoculation with the *I. marina* consortium. **C.** Images of marine broth plates inoculated with samples taken of *I. marina* cultures treated with antibiotic cocktails compared to control.

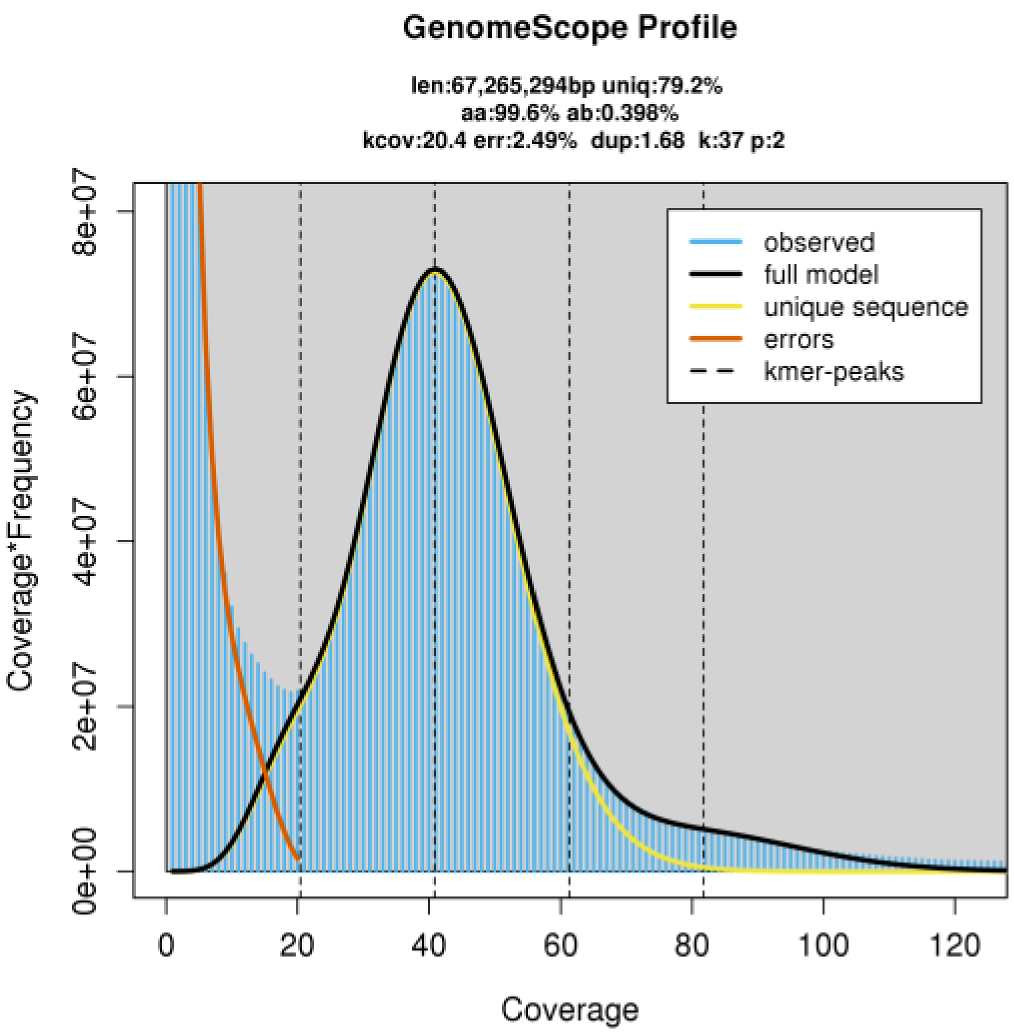


GenomeScope version 2.0

p = 2
k = 37

| property | min | max |
|---|---|---|
| Homozygous (aa) | 99.5901% | 99.6141% |
| Heterozygous (ab) | 0.3859% | 0.40985% |
| Genome Haploid Length | 67,125,052 bp | 67,265,294 bp |
| Genome Repeat Length | 13,953,246 bp | 13,982,398 bp |
| Genome Unique Length | 53,171,806 bp | 53,282,896 bp |
| Model Fit | 85.3674% | 99.0997% |
| Read Error Rate | 2.48741% | 2.48741% |

**Figure S3. Plot of k-mer frequencies in the reads used for the *Incisomonas marina* MAG**. K-mer size of 38, as predicted using Genomescope2 (Materials and Methods). The single peak demonstrates the haploid nature of the genome. Plot generated by Genomescope2 (Ranallo-Benavidez et al. 2020).

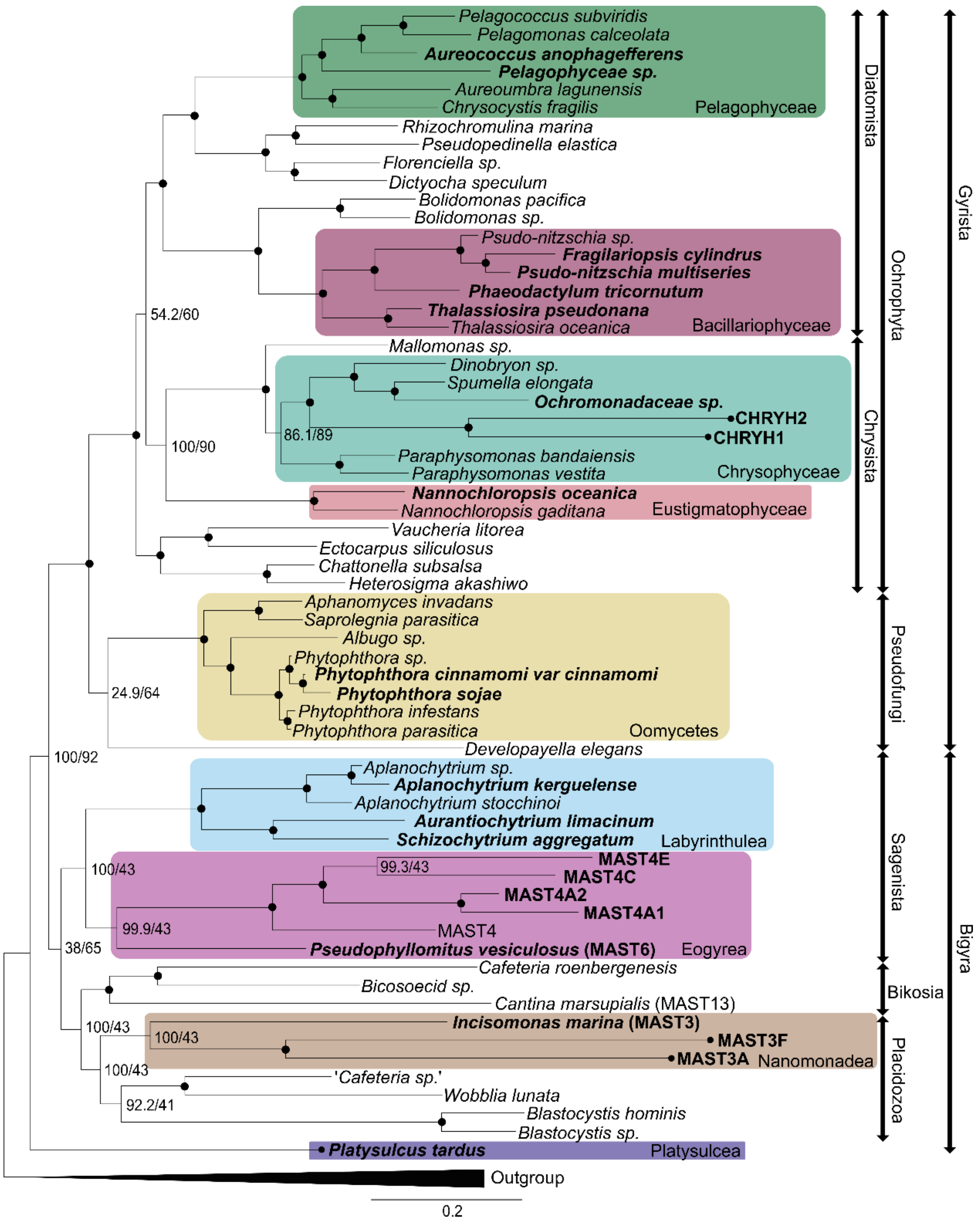


**Figure S4. Phylogenomic tree of the Stramenopila.** The tree was created using a concatenated alignment of 120 genes with columns removed where 20% of sites were gaps. Gene alignments from Thakur et al., (2019) (Thakur et al. 2019) were used with additional species of interest included to increase taxon sampling. IQtree was used in model prediction mode to create a maximum likelihood tree using ultrafast bootstrapping and the -altr flag, resulting in two values of node support. The model used was VT+F+I+G4. Taxonomic ranking on the right of the tree reflects those of Thakur et al., 2019 (Thakur et al. 2019). The outgroup includes species from the Rhizaria and Alveolata. Nodes with support values 100/100 are indicated with a filled black circle.

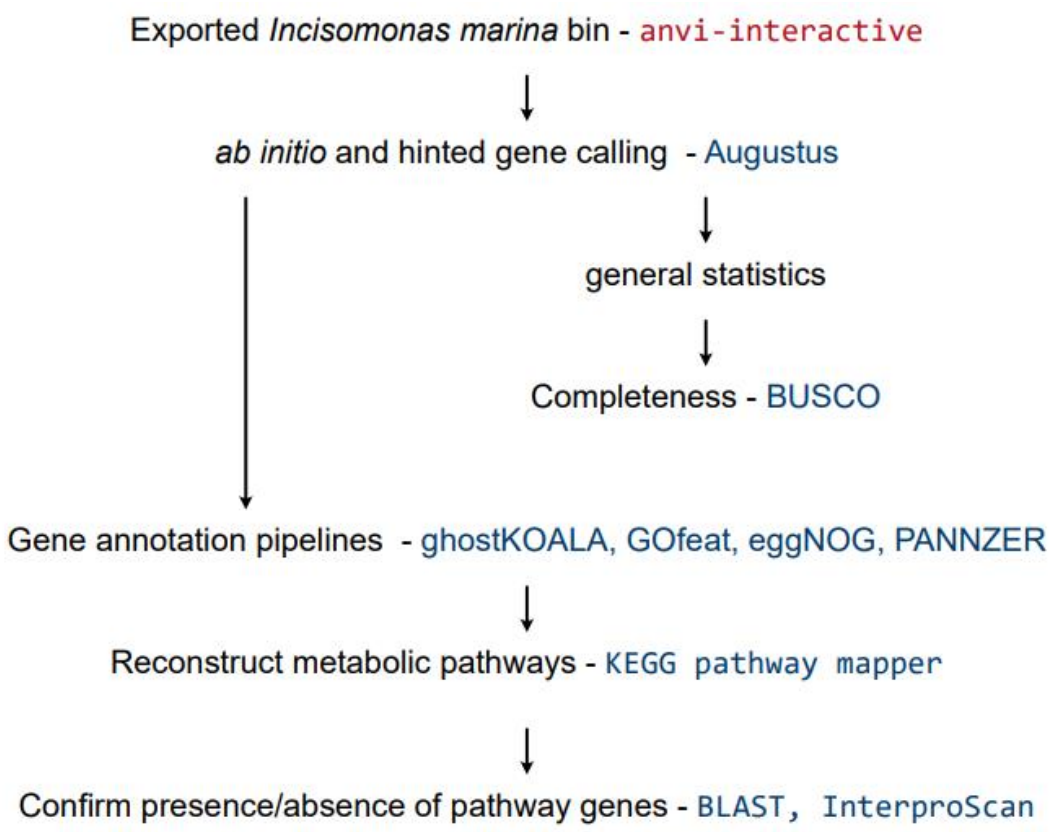


**Figure S5. Schematic of the downstream analysis of the *Incisomonas marina* MAG**. This includes gene calling, completeness estimation, annotation and pathway analysis.

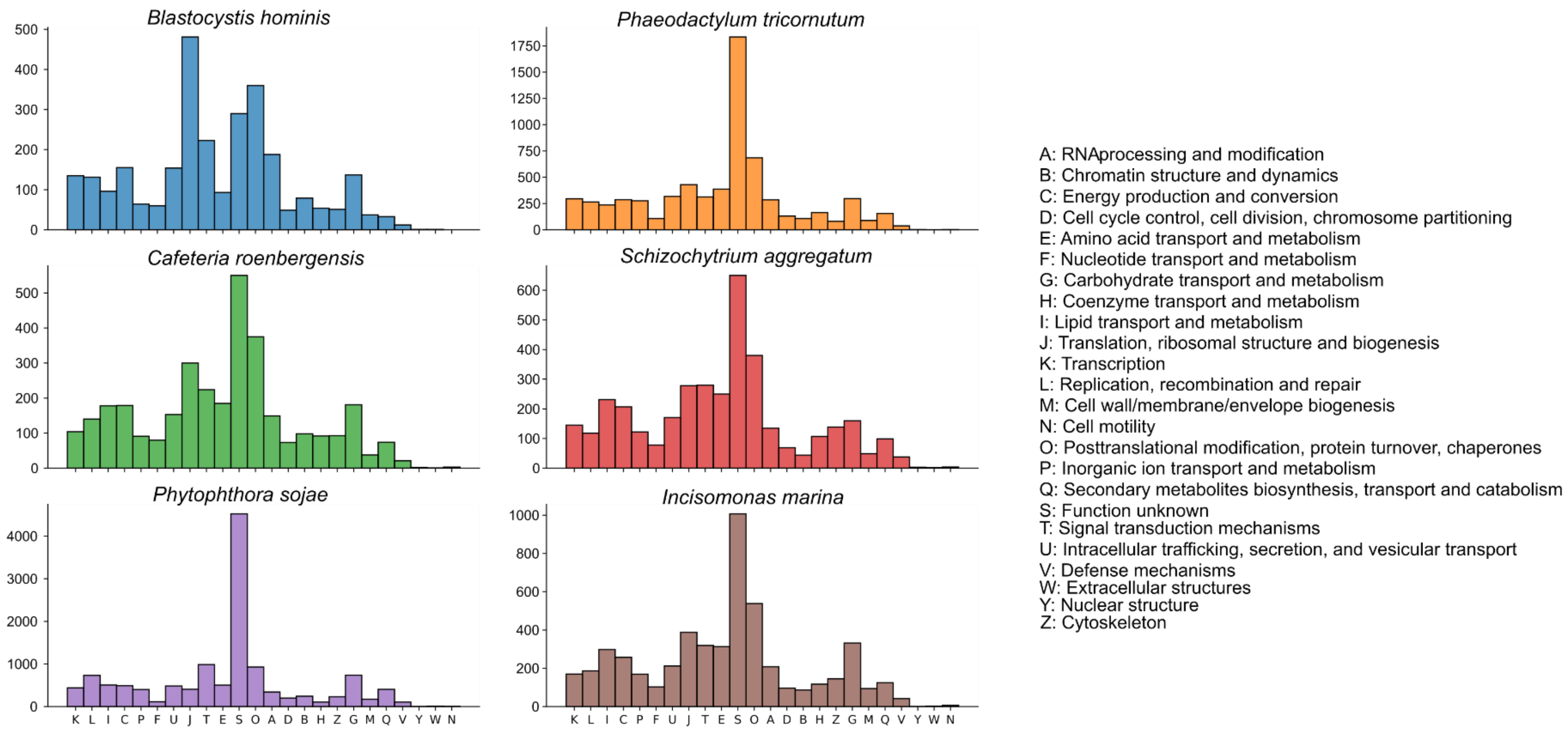


**Figures S6**. **Comparison of the percentage of Clusters of Orthologous Genes (COG) category assignments across the annotated genes of six stramenopiles**. Stramenopile species used for this comparison included *Incisomonas marina* from this study, *Cafeteria roenbergensis*, *Blastocystis hominis*, *Phaeodactylum tricornutum*, *Phytophthora sojae* and *Schizochytrium aggregatum*. Each organism displays a relatively similar functional landscape at this macro-scale of annotation.

**Table S1. Summary of antibiotic cocktails used to treat *Incisomonas marina* cultures.** Concentrations successfully employed in the literature to treat cultures of other stramenopile taxa are also given: [a](Wilkens and Maas 2012); [b](Droop 1967); [c](Ferrante et al. 2020); [d](Koedooder et al. 2019).

| **Antibiotic** | **Cocktail 1** (µg/ml) (this study) | **Cocktail 2** (µg/ml) (this study) | **Concentrations used in the literature against other Stramenopile taxa** (µg/ml) | **Taxa** |
|---|---|---|---|---|
| Rifampicin | 10 | - | 300[a] | Thraustrochytrids |
| Streptomycin | 100 | 30 | 1600[b] | Diatoms |
| Ampicillin | 100 | 50 | 700[c] | Diatoms |
| Gentamycin | 100 | - | 50-100[d] | Diatoms |
| Neomycin | - | 60 | 400[b] | Diatoms |
| Kanamycin | - | 50 | >250[d] | Diatoms |
| Chloramphenicol | - | 25 | 400[b] | Diatoms |

**Table S2. Summary statistics (1 of 2) of metagenome bins of *Incisomonas marina* and bacterial isolates.** The metagenome was assembled using OxyNanopore long-read data and polished using Derrelle et al., (2016) Illumina short-read data (Derelle et al. 2016) with Flye and Pilon respectively. Supervised binning was performed using the Anvi'o interactive interface (Eren et al. 2015).

| *Bin No.* | Variability | Relative abundance | Std coverage | Mean coverage Q2Q3 | Mean coverage | Max normalised ratio | Abundance | Bins % recruitment | Detection % |
|---|---|---|---|---|---|---|---|---|---|
| *1* | 51.26 | 1 | 8.88 | 85.66 | 85.48 | 1 | 0.90 | 36.83 | 99.72 |
| *2* | 50.00 | 1 | 30.61 | 426.92 | 424.67 | 1 | 4.49 | 11.18 | 99.99 |
| *3* | 32.73 | 1 | 5.27 | 57.19 | 57.14 | 1 | 0.60 | 1.65 | 100.00 |
| *4* | 2.05 | 1 | 2.83 | 19.06 | 19.07 | 1 | 0.20 | 0.35 | 100.00 |
| *5* | 27.53 | 1 | 18.68 | 193.69 | 192.72 | 1 | 2.04 | 3.49 | 99.61 |
| *6* | 26.08 | 1 | 9.94 | 153.42 | 152.93 | 1 | 1.62 | 3.79 | 100.00 |
| *7* | 0.14 | 1 | 2.48 | 4.80 | 5.15 | 1 | 0.05 | 0.10 | 99.85 |
| *8* | 35.68 | 1 | 5.52 | 61.74 | 61.68 | 1 | 0.65 | 2.16 | 100.00 |
| *9* | 28.52 | 1 | 24.68 | 554.37 | 551.61 | 1 | 5.83 | 13.94 | 100.00 |
| *10* | 0.60 | 1 | 2.02 | 9.32 | 9.36 | 1 | 0.10 | 0.25 | 100.00 |
| *11* | 18.33 | 1 | 3.93 | 35.26 | 35.25 | 1 | 0.37 | 0.80 | 100.00 |
| *12* | 44.31 | 1 | 9.59 | 150.10 | 149.88 | 1 | 1.58 | 3.55 | 99.82 |
| *13* | 45.63 | 1 | 6.62 | 87.08 | 86.96 | 1 | 0.92 | 2.28 | 100.00 |
| *14* | 7.19 | 1 | 3.33 | 24.37 | 24.33 | 1 | 0.26 | 0.72 | 100.00 |
| *15* | 48.14 | 1 | 13.63 | 234.79 | 234.19 | 1 | 2.47 | 6.64 | 100.00 |
| *16* | 51.77 | 1 | 7.06 | 86.61 | 86.54 | 1 | 0.91 | 2.78 | 100.00 |
| *17* | 0.55 | 1 | 30.05 | 6.88 | 16.83 | 1 | 0.18 | 0.35 | 99.92 |
| *18* | 131.61 | 1 | 7.72 | 74.34 | 73.84 | 1 | 0.78 | 2.39 | 100.00 |
| *19* | 6.92 | 1 | 2.64 | 18.50 | 18.53 | 1 | 0.20 | 0.42 | 100.00 |
| *20* | 45.23 | 1 | 8.49 | 130.26 | 129.93 | 1 | 1.37 | 3.97 | 100.00 |
| *21* | 2.37 | 1 | 2.09 | 9.48 | 9.53 | 1 | 0.10 | 0.18 | 99.91 |
| *22* | 0.64 | 1 | 1.75 | 6.58 | 6.60 | 1 | 0.07 | 0.18 | 99.91 |
| *23* | 3.61 | 1 | 2.38 | 13.98 | 14.02 | 1 | 0.15 | 0.39 | 99.99 |
| *24* | 1.79 | 1 | 2.52 | 14.57 | 14.59 | 1 | 0.15 | 0.35 | 100.00 |
| *Un-binned* | - | - | - | - | - | - | - | 1.27 | - |

**Table S3. Summary statistics (2 of 2) of metagenome bins of *Incisomonas marina* and bacterial isolates**. The metagenome was assembled using OxyNanopore long-read data and polished using Derrelle et al., (2016) Illumina short-read data (Derelle et al. 2016) with Flye and Pilon respectively. Supervised binning was performed using the Anvi'o interactive interface (Eren et al. 2015). *Percent completion for bin 1 using the standard Anvi'o eukaryotic BUSCO library. However, when the stramenopile library was used, completion was 93.0%

| Bin No. | Total length (bp) | Number contigs | N50 | GC content | Percent completion | Percent redundancy |
|---|---|---|---|---|---|---|
| 1 | 68,357,064 | 213 | 1215950 | 52.5 | 68.7* | 16.9 |
| 2 | 4,108,619 | 5 | 4006367 | 40.9 | 100 | 5.6 |
| 3 | 4,510,811 | 3 | 4487859 | 43.3 | 100 | 2.8 |
| 4 | 2,913,340 | 1 | 2913340 | 44.8 | 98.6 | 4.2 |
| 5 | 2,900,267 | 63 | 94070 | 39.4 | 0 | 0 |
| 6 | 3,899,295 | 2 | 3863077 | 33.4 | 97.2 | 16.9 |
| 7 | 3,085,312 | 21 | 266584 | 39.0 | 78.9 | 15.5 |
| 8 | 5,505,433 | 2 | 5469367 | 39.7 | 98.6 | 2.8 |
| 9 | 3,945,902 | 1 | 3945902 | 41.6 | 98.6 | 4.2 |
| 10 | 4,219,735 | 4 | 2482457 | 49.6 | 95.8 | 7.0 |
| 11 | 3,566,829 | 1 | 3566829 | 63.6 | 97.2 | 2.8 |
| 12 | 3,706,920 | 3 | 3586063 | 57.0 | 100 | 2.8 |
| 13 | 4,096,611 | 1 | 4096611 | 57.4 | 100 | 0 |
| 14 | 4,658,551 | 1 | 4658551 | 61.5 | 100 | 0 |
| 15 | 4,425,738 | 1 | 4425738 | 61.0 | 97.2 | 8.5 |
| 16 | 5,035,835 | 4 | 4700859 | 61.7 | 97.2 | 1.4 |
| 17 | 4,175,398 | 17 | 605885 | 60.5 | 90.1 | 9.9 |
| 18 | 5,058,777 | 2 | 4789643 | 59.1 | 98.6 | 0 |
| 19 | 3,516,122 | 1 | 3516122 | 58.0 | 97.2 | 7.0 |
| 20 | 4,785,063 | 2 | 4571768 | 53.5 | 97.2 | 0 |
| 21 | 2,943,871 | 8 | 842461 | 46.9 | 93.0 | 5.6 |
| 22 | 4,331,361 | 10 | 543885 | 44.9 | 98.6 | 9.9 |
| 23 | 4,324,906 | 2 | 4265747 | 66.8 | 97.2 | 7.0 |
| 24 | 3,704,244 | 1 | 3704244 | 62.7 | 100 | 0 |

**Table S4. Taxonomic predictions of each bin as predicted by Anvi'o taxonomic prediction.** There is no taxonomic prediction for bin 1 as this is the eukaryotic bin belonging to *I. marina,* indicated by the identified locations of 18S/28S rRNA genes.

| *Bin No.* | Domain | Phylum | Class | Order | Family | Genus | Species |
|---|---|---|---|---|---|---|---|
| *1* | - | - | - | - | - | - | - |
| *2* | *Bacteria* | *Proteobacteria* | *Gammaproteobacteria* | *Enterobacterales* | *Alteromonadaceae* | *Pseudoalteromonas* | - |
| *3* | *Bacteria* | *Proteobacteria* | *Gammaproteobacteria* | *Enterobacterales* | *Alteromonadaceae* | *Alteromonas* | - |
| *4* | *Bacteria* | *Proteobacteria* | *Gammaproteobacteria* | *Nitrosococcales* | *Methylophagaceae* | *Methylophaga* | - |
| *5* | - | - | - | - | - | - | - |
| *6* | *Bacteria* | *Bacteroidota* | *Bacteroidia* | *Flavobacteriales* | *Flavobacteriaceae* | *Winogradskyella* | ***Marinobacter hydrocarbonoclasticus **** |
| *7* | *Bacteria* | *Bacteroidota* | *Bacteroidia* | *Flavobacteriales* | *Crocinitomicaceae* | - | - |
| *8* | *Bacteria* | *Bacteroidota* | *Bacteroidia* | *Flavobacteriales* | *Flavobacteriaceae* | *Arenibacter* | - |
| *9* | *Bacteria* | *Bacteroidota* | *Bacteroidia* | *Flavobacteriales* | *Flavobacteriaceae* | *Muricauda* | - |
| *10* | *Bacteria* | *Proteobacteria* | *Gammaproteobacteria* | *Pseudomonadales* | *Spongiibacteraceae* | *Zhongshania* | *Zhongshania sp002915595* |
| *11* | *Bacteria* | *Proteobacteria* | *Alphaproteobacteria* | *Rhodobacterales* | *Rhodobacteraceae* | *Roseovarius* | *Roseovarius halotolerans* |
| *12* | *Bacteria* | *Proteobacteria* | *Alphaproteobacteria* | *Rhodobacterales* | *Rhodobacteraceae* | *Sulfitobacter* | *Sulfitobacter marinus* |
| *13* | *Bacteria* | *Proteobacteria* | *Gammaproteobacteria* | *Pseudomonadales* | *Oleiphilaceae* | *Marinobacter* | *Marinobacter hydrocarbonoclasticus** |
| *14* | *Bacteria* | *Proteobacteria* | *Gammaproteobacteria* | *Pseudomonadales* | *Alcanivoracaceae* | *Alcanivorax* | - |
| *15* | *Bacteria* | *Proteobacteria* | *Alphaproteobacteria* | *Rhodobacterales* | *Rhodobacteraceae* | *Epibacterium* | *Epibacterium scottomollicae* |
| *16* | *Bacteria* | *Proteobacteria* | *Alphaproteobacteria* | *Rhodobacterales* | *Rhodobacteraceae* | *Antarctobacter* | *Antarctobacter heliothermus* |
| *17* | *Bacteria* | *Proteobacteria* | *Alphaproteobacteria* | *Rhizobiales* | *Rhizobiaceae* | *Pararhizobium* | - |
| *18* | *Bacteria* | *Proteobacteria* | *Alphaproteobacteria* | *Rhizobiales* | *Rhizobiaceae* | *Hoeflea* | - |
| *19* | *Bacteria* | *Proteobacteria* | *Alphaproteobacteria* | *Caulobacterales* | *Hyphomonadaceae* | *Hyphomonas* | *Hyphomonas atlantica* |
| *20* | *Bacteria* | *Proteobacteria* | *Alphaproteobacteria* | *Rhodospirillales* | *Thalassospiraceae* | *Thalassospira* | - |
| *21* | *Bacteria* | *Proteobacteria* | *Gammaproteobacteria* | *Enterobacterales* | *Alteromonadaceae* | *Idiomarina* | *Idiomarina loihiensis* |
| *22* | *Bacteria* | *Proteobacteria* | *Gammaproteobacteria* | *Pseudomonadales* | *Marinomonadaceae* | *Marinomonas* | *Marinomonas sp004352855* |
| *23* | *Bacteria* | *Proteobacteria* | *Alphaproteobacteria* | *Rhodobacterales* | *Rhodobacteraceae* | *Pseudooceanicola* | *Pseudooceanicola marinus** |
| *24* | *Bacteria* | *Proteobacteria* | *Alphaproteobacteria* | *Parvibaculales* | *Parvibaculaceae* | - | - |

**Table S5**. Red algal annotated genes in *Incisomonas marina*, as annotated by ghostKOALA (Kanehisa et al. 2015). Evidence for plastid associated red algal genes were derived from https://www.arabidopsis.org/servlets/TairObject?type=locus&name=AT1G80380. Predicted subcellular localisation of corresponding *I. marina* genes as inferred by HECTAR (Gschloessl et al. 2008) are also given. Key: n = not predicted.

| Gene ID | KO number | Description | Predicted Cp/Mt localisation of *I. marina* predicted protein via HECTAR | Plastid linked? |
|---|---|---|---|---|
| **g44.t1** | K08288 | PRKCSH; protein kinase C substrate 80K-H | n | n |
| **g504.t1** | K19612 | PDE12; 2',5'-phosphodiesterase [EC:3.1.13.4 3.1.4.-] | n | Mitochondrial |
| **g569.t1** | K18156 | ATP23, XRCC6BP1; mitochondrial inner membrane protease ATP23 [EC:3.4.24.-] | n | Mitochondrial |
| **g1394.t1** | K01875 | SARS, serS; seryl-tRNA synthetase [EC:6.1.1.11] | n | Mitochondrial |
| **g1556.t1** | K00942 | gmk, GUK1; guanylate kinase [EC:2.7.4.8] | n | n |
| **g1679.t1** | K05917 | CYP51; sterol 14alpha-demethylase [EC:1.14.14.154 1.14.15.36] | n | n |
| **g1705.t1** | K07305 | msrB; peptide-methionine (R)-S-oxide reductase [EC:1.8.4.12] | n | n |
| **g1872.t1** | K18156 | ATP23, XRCC6BP1; mitochondrial inner membrane protease ATP23 [EC:3.4.24.-] | n | Mitochondrial |
| **g1888.t1** | K00939 | adk, AK; adenylate kinase [EC:2.7.4.3] | n | n |
| **g1969.t1** | K08592 | SENP1; sentrin-specific protease 1 [EC:3.4.22.68] | n | n |
| **g2756.t1** | K20347 | TMED2, EMP24; p24 family protein beta-1 | n | n |
| **g3076.t1** | K12821 | PRPF40, PRP40; pre-mRNA-processing factor 40 | n | n |
| **g3633.t1** | K08288 | PRKCSH; protein kinase C substrate 80K-H | n | n |
| **g3942.t1** | K08592 | SENP1; sentrin-specific protease 1 [EC:3.4.22.68] | n | n |
| **g4827.t1** | K11498 | CENPE; centromeric protein E | n | n |
| **g4368.t1** | K08796 | BRSK; BR serine/threonine kinase [EC:2.7.11.1] | n | n |
| **g4619.t1** | K00939 | adk, AK; adenylate kinase [EC:2.7.4.3] | n | n |
| **g4872.t1** | K11498 | CENPE; centromeric protein E | n | n |
| **g4885.t1** | K08592 | SENP1; sentrin-specific protease 1 [EC:3.4.22.68] | n | n |
| **g5477.t1** | K11498 | CENPE; centromeric protein E | n | n |
| **g5867.t1** | K00939 | adk, AK; adenylate kinase [EC:2.7.4.3] | n | n |
| **g6309.t1** | K08288 | PRKCSH; protein kinase C substrate 80K-H | n | n |
| **g7000.t1** | K20347 | TMED2, EMP24; p24 family protein beta-1 | n | n |
| **g7257.t1** | K00939 | adk, AK; adenylate kinase [EC:2.7.4.3] | n | n |

| | | | | |
|---|---|---|---|---|
| **g7265.t1** | K13703 | ABHD11; abhydrolase domain-containing protein 11 | Mitochondrial | n |
| **g8047.t1** | K12821 | PRPF40, PRP40; pre-mRNA-processing factor 40 | n | n |
| **g8708.t1** | K08592 | SENP1; sentrin-specific protease 1 [EC:3.4.22.68] | n | n |
| **g9448.t1** | K00939 | adk, AK; adenylate kinase [EC:2.7.4.3] | n | n |
| **g9925.t1** | K08592 | SENP1; sentrin-specific protease 1 [EC:3.4.22.68] | n | n |
| **g10230.t1** | K17777 | TIM9; mitochondrial import inner membrane translocase subunit TIM9 | n | Mitochondrial |
| **g10403.t1** | K12821 | PRPF40, PRP40; pre-mRNA-processing factor 40 | n | n |
| **g10658.t1** | K00939 | adk, AK; adenylate kinase [EC:2.7.4.3] | n | n |
| **g11052.t1** | K03469 | rnhA, RNASEH1; ribonuclease HI [EC:3.1.26.4] | n | n |
| **g11660.t1** | K12821 | PRPF40, PRP40; pre-mRNA-processing factor 40 | n | n |
| **g11580.t1** | K00939 | adk, AK; adenylate kinase [EC:2.7.4.3] | n | n |
| **g11705.t1** | K08592 | SENP1; sentrin-specific protease 1 [EC:3.4.22.68] | n | n |
| **g12317.t1** | K18703 | SUGCT; succinate---hydroxymethylglutarate CoA-transferase [EC:2.8.3.13] | n | Mitochondrial |
| **g13694.t1** | K11498 | CENPE; centromeric protein E | n | n |
| **g16639.t1** | K01875 | SARS, serS; seryl-tRNA synthetase [EC:6.1.1.11] | n | Mitochondrial |
| **g12792.t1** | K08592 | SENP1; sentrin-specific protease 1 [EC:3.4.22.68] | n | n |
| **g12837.t1** | K12821 | PRPF40, PRP40; pre-mRNA-processing factor 40 | n | n |
| **g13091.t1** | K24887 | GTPBP1; GTP-binding protein 1 | n | n |
| **g13095.t1** | K00939 | adk, AK; adenylate kinase [EC:2.7.4.3] | n | n |
| **g13290.t1** | K06675 | SMC4; structural maintenance of chromosome 4 | n | n |
| **g13486.t1** | K08592 | SENP1; sentrin-specific protease 1 [EC:3.4.22.68] | n | n |
| **g13520.t1** | K12821 | PRPF40, PRP40; pre-mRNA-processing factor 40 | n | n |
| **g13581.t1** | K15111 | SLC25A26; solute carrier family 25 (mitochondrial S-adenosylmethionine transporter), member 26 | n | Mitochondrial |
| **g13694.t1** | K11498 | CENPE; centromeric protein E | n | n |
| **g13783.t1** | K12812 | DDX39B, UAP56, SUB2; ATP-dependent RNA helicase UAP56/SUB2 [EC:3.6.4.13] | n | n |
| **g13784.t1** | K12812 | DDX39B, UAP56, SUB2; ATP-dependent RNA helicase UAP56/SUB2 [EC:3.6.4.13] | n | n |
| **g13943.t1** | K01726 | GAMMACA; gamma-carbonic anhydrase [EC:4.2.1.-] | n | Mitochondrial |
| **g13948.t1** | K08288 | PRKCSH; protein kinase C substrate 80K-H | n | n |
| **g14169.t1** | K11090 | LA, SSB; lupus La protein | n | n |
| **g14773.t1** | K11498 | CENPE; centromeric protein E | n | n |
| **g15451.t1** | K00942 | gmk, GUK1; guanylate kinase [EC:2.7.4.8] | n | n |
| **g15645.t1** | K06002 | PGA; pepsin A [EC:3.4.23.1] | n | n |
| **g16100.t1** | K08288 | PRKCSH; protein kinase C substrate 80K-H | n | n |

| | | | | |
|---|---|---|---|---|
| **g16227.t1** | K08592 | SENP1; sentrin-specific protease 1 [EC:3.4.22.68] | n | n |
| **g16320.t1** | K18932 | ZDHHC; palmitoyltransferase [EC:2.3.1.225] | n | n |
| **g16412.t1** | K03469 | rnhA, RNASEH1; ribonuclease HI [EC:3.1.26.4] | n | n |
| **g16639.t1** | K01875 | SARS, serS; seryl-tRNA synthetase [EC:6.1.1.11] | Mitochondrial | n |
| **g17092.t1** | K15918 | GLYK; D-glycerate 3-kinase [EC:2.7.1.31] | n | Chloroplast, mitochondria, nucleus |
| **g17171.t1** | K15111 | SLC25A26; solute carrier family 25 (mitochondrial S-adenosylmethionine transporter), member 26 | n | n |
| **g17341.t1** | K08592 | SENP1; sentrin-specific protease 1 [EC:3.4.22.68] | n | n |
| **g17981.t1** | K05906 | PCYOX1, FCLY; prenylcysteine oxidase / farnesylcysteine lyase [EC:1.8.3.5 1.8.3.6] | n | n |
| **g18240.t1** | K11498 | CENPE; centromeric protein E | n | n |
| **g18296.t1** | K18932 | ZDHHC; palmitoyltransferase [EC:2.3.1.225] | n | n |
| **g18501.t1** | K18932 | ZDHHC; palmitoyltransferase [EC:2.3.1.225] | n | n |
| **g17341.t1** | K08592 | SENP1; sentrin-specific protease 1 [EC:3.4.22.68] | n | n |
| **g18514.t1** | K08592 | SENP1; sentrin-specific protease 1 [EC:3.4.22.68] | n | n |
| **g18609.t1** | K08592 | SENP1; sentrin-specific protease 1 [EC:3.4.22.68] | n | n |
| **g19413.t1** | K07305 | msrB; peptide-methionine (R)-S-oxide reductase [EC:1.8.4.12] | Mitochondrial | n |
| **g19479.t1** | K01726 | GAMMACA; gamma-carbonic anhydrase [EC:4.2.1.-] | n | n |
| **g19493.t1** | K08288 | PRKCSH; protein kinase C substrate 80K-H | n | n |

**Table S6. Presence/absence of vitamin C biosynthesis enzymes L-galactonolactone dehydrogenase (GLDH) and L-gulonolactone oxidase (GULO) in the genomes of six sequenced stramenopiles.** Protein identifiers are also given for each hit.

| **Species** | **GLDH** | **GULO** |
|---|---|---|
| *Incisomonas marina* | g8348.t1 | g11511.t1 |
| *Schizochytrium aggregatum* | 98185 | - |
| *Phytophthora sojae* | 485337 | - |
| *Phaeodactylum tricornutum* | 23292 | - |
| *Blastocystis hominis* Singapore Isolate B | - | - |
| *Cafeteria roenbergensis* BVI | 622 | 7535 |

**Table S7**. Description of hits retrieved from the *Tara Oceans* Eukaryotic metagenome and single-cell assembled genome database (EUK_SMAGs) when blasting the *Incisomonas marina* DSYB predicted protein sequence, with an e-value cut-off of $1e \times 10^{-70}$. The taxonomic classification of each MAG, metabolic mode and BUSCO completion as assigned by (Delmont et al. 2022) is also given, with MAST taxa shaded in grey. Predicted subcellular targeting by HECTAR (Gschloessl et al. 2008) is also given.

| Protein hit Id. | MAG Id. | MAG Taxonomic Assignment by Delmont et al., 2022 | Metabolic mode | BUSCO Completion (%) | Predicted targeting |
|---|---|---|---|---|---|
| SMAGs_714947 | TARA_AON_82_MAG_00276_000000004952.4.1 | MAST-7 | Heterotrophic | 30.2 | Mitochondrial |
| SMAGs_9697997 | TARA_ARC_108_MAG_00241_000000000996.1.1 | MAST unclassified | Heterotrophic | 38.8 | Mitochondrial |
| SMAGs_2685109 | TARA_ARC_108_MAG_00241_000000000996.1.1 | MAST unclassified | Heterotrophic | 38.8 | Other |
| SMAGs_5412876 | TARA_MED_95_MAG_00408_000000009893.2.1 | New Oomycota | Heterotrophic | 53.8 | Other |
| SMAGs_5409111 | TARA_MED_95_MAG_00408_000000009893.2.1 | New Oomycota | Heterotrophic | 53.8 | Other |
| SMAGs_4706793 | TARA_IOS_50_MAG_00124_000000001544.5.1 | Aureococcus | Photosynthetic | 54.9 | Other |
| SMAGs_1750378 | TARA_AOS_82_MAG_00144_000000003998.3.1 | New_Chrysochromulinaceae_01 | Photosynthetic | 27.1 | Mitochondrial |
| SMAGs_4614449 | TARA_ION_45_MAG_00201_000000002677.5.1 | New_Chrysochromulinaceae_01 | Photosynthetic | 74.9 | Mitochondrial |
| SMAGs_6824365 | TARA_PON_109_MAG_00232_000000000346.1.1 | New_Chrysochromulinaceae_01 | Photosynthetic | 52.2 | Mitochondrial |
| SMAGs_8443951 | TARA_PSW_86_MAG_00280_000000000285.16.1 | New_Chrysochromulinaceae_01 | Photosynthetic | 32.2 | Mitochondrial |
| SMAGs_4027617 | TARA_ARC_108_MAG_00326_000000006951.13.4 | New_Chrysochromulinaceae_01 | Photosynthetic | 68.6 | Mitochondrial |
| SMAGs_8506517 | TARA_PSW_86_MAG_00285_000000000349.2.1 | New_Chrysochromulinaceae_01 | Photosynthetic | 31.8 | Signal peptide |
| SMAGs_720636 | TARA_AON_82_MAG_00278_000000002804.3.1 | New_Chrysochromulinaceae_01 | Photosynthetic | 27.8 | Signal peptide |
| SMAGs_3784438 | TARA_ARC_108_MAG_00319_000000002099.3.1 | Phaeocystis | Photosynthetic | 56.9 | Mitochondrial |
| SMAGs_10058803 | TARA_SOC_28_MAG_00074_000000006525.6.2 | Phaeocystis | Photosynthetic | 52.6 | Mitochondrial |
| SMAGs_4145110 | TARA_ION_45_MAG_00152_000000003198.10.1 | Sister_Phaeocystis | Photosynthetic | 43.6 | Signal peptide |
| SMAGs_5732555 | TARA_MED_95_MAG_00437_000000002734.2.2 | Picozoa | Heterotrophic | 21.6 | Mitochondrial |
| SMAGs_3725435 | TARA_ARC_108_MAG_00317_000000000411.3.1 | Picozoa | Heterotrophic | 75.7 | Other |
| SMAGs_10171271 | TARA_SOC_28_MAG_00079_000000015284.10.1 | Picozoa | Heterotrophic | 75.7 | Mitochondrial |
| SMAGs_8949003 | TARA_PSW_86_MAG_00314_000000005600.2.1 | Picozoa | Heterotrophic | 62.0 | Mitochondrial |
| SMAGs_10199646 | TARA_SOC_28_MAG_00079_000000002210.2.1 | Picozoa | Heterotrophic | 75.7 | Other |
| SMAGs_2909318 | TARA_ARC_108_MAG_00257_000000000227.17.2 | Chonoflagellatea_NA | Heterotrophic | 25.1 | Mitochondrial |
| SMAGs_2830157 | TARA_ARC_108_MAG_00250_000000008414.1.2 | Chonoflagellatea_NA | Heterotrophic | 40.4 | Mitochondrial |

| | | | | | |
|---|---|---|---|---|---|
| SMAGs_5447382 | TARA_MED_95_MAG_00409_000000011530.1.1 | Ochrophyta_Dictyochophyceae_NA | Photosynthetic | 72.6 | Other |
